# Oncofetal RNA-binding proteins of IGF2BP family suppress IRF-3 and NF-kB dependent transcription downstream of cytosolic RNA sensors

**DOI:** 10.64898/2026.05.22.726816

**Authors:** Garrett Knox, Ruba Agili-Shaban, Alexis Morrissey, Karamveer Karamveer, Ahsan Polash, Maria Wohlbowne, Stefanija Kinzy, Cheryl Keller, Todd Schell, Jeremy Hengst, George-Lucian Moldovan, Junjia Zhu, Arati Sharma, Hong Zheng, Edward Harhaj, Markus Hafner, Zhenqiu Liu, Yasin Uzun, Shaun Mahony, Irina Elcheva

## Abstract

Activation of innate inflammatory signaling and tumor-specific antigen presentation in cancer cells provides a foundation for anti-cancer immunotherapies. Here, we show that the insulin-like growth factor 2 mRNA-binding proteins (IGF2BP1, IGF2BP2, and IGF2BP3), which are upregulated across diverse human malignancies, including acute myeloid leukemia (AML), suppress RNA-sensing pattern recognition receptor signaling and downstream ISRE-and NF-κB-dependent transcription. Among these pathways, RIG-I signaling is particularly sensitive to IGF2BP-mediated inhibition. This suppressive effect is strongest when all three IGF2BP paralogs are co-expressed, especially in embryonic-like hematoendothelial cells and leukemia stem cells. Mechanistically, IGF2BPs suppress innate immune signaling by directly binding to TNFAIP3 mRNA and promoting its expression. Consequently, inhibition of IGF2BPs activates innate immune signaling and induces MHC class I gene expression in AML cells, highlighting IGF2BPs as promising therapeutic targets to enhance RIG-I- and TLR-based cancer immunotherapies.

**Highlights:**

- IGF2BP family of proteins regulate PRR responses
- IGF2BP levels affect chromatin accessibility and pan-transcriptome
- SV40 large T antigen induces *de novo* expression of IGF2BPs
- TNFAIP3 mRNA levels and stability are regulated by IGF2BPs

## Introduction

RNA-binding proteins (RBPs) control all steps of RNA fate from transcription to translation and decay. RBPs play a critical role in embryonic development whereas their malfunction, due to mutations or aberrant expression, is associated with various diseases including cancer (Elcheva and Spiegelman, 2021; Hentze et al., 2018). Insulin-like growth factor 2 mRNA binding proteins (human paralogs IGF2BP1, IGF2BP2, and IGF2BP3) are a family of cytoplasmic proteins largely known as positive regulators of mRNA stability and translation in mammalian cells (Degrauwe et al., 2016). Normally expressed during embryonic development, IGF2BP1 and IGF2BP3 are silenced in most adult tissues, while basal expression of IGF2BP2 remains through adulthood. Among direct RNA targets of IGF2BPs are transcription factors, cytoskeletal proteins, cellular receptors of growth and proliferation, regulators of metabolism (e.g., *c-MYC, ACTB, CD44, VEGFA, MYB, HOXB4, SLC2A1, ALDHA1A1)(Doyle et al., 1998; Elcheva et al., 2020a; Stöhr et al., 2012).* Critical supporters of G1/S cycle progression, IGF2BPs are upregulated in 30-70% of cancers depending on histological type and are linked to worse overall survival (Huang et al., 2018; Köbel et al., 2007; Palanichamy et al., 2016; Weidensdorfer et al., 2009).

Recently, we discovered that in addition to cancer cell-autonomous pro-proliferative role, IGF2BPs function as negative regulators of interferon-stimulated genes (ISGs) expression (Elcheva et al., 2023). We showed that genetic inhibition of IGF2BPs significantly increases expression of interferon type I/II genes in normal and malignant mammalian cells, enhances infiltration of myeloid and NKT cells into tumor microenvironment and improves outcomes of anti-PD-1 therapies in mouse model of melanoma. Other reports support observation about immunosuppressive properties of IGF2BPs in cancer cells, however, cellular and molecular mechanisms of their effect on immune responses are unknown (Bley et al., 2025; Peng et al., 2024; Tang et al., 2024).

In this study, we show that inhibition of IGF2BPs enhances ISRE-and NF-κB-dependent transcription from corresponding ISRE-and NF-κB-controlled promoters, both with and without cytokine stimulation. The endogenous priming of proinflammatory signaling in IGF2BP1–3 knockout (KO) cells is accompanied by significant upregulation of RNA-sensing pattern recognition receptors (PRRs) and antigen presentation (MHC class I) proteins, which is further enhanced upon treatment with RIG-I, TLR3, and TLR8 agonists. Here, we employed cellular systems co-expressing three IGF2BP paralogs which allow for a comparative analysis of their functions: embryonic-like human induced pluripotent stem cells (hiPSC) and their derivatives, Simian Virus 40 (SV40) large T antigen expressing mammalian cells, and acute myeloid leukemia (AML) with various patterns of IGF2BPs expression. We found that the IGF2BP-mediated immunosuppressive phenotype is more pronounced when the embryonic-specific paralogs IGF2BP1 and IGF2BP3 are co-expressed, suggesting that acquisition of a stem cell-like phenotype reduces intracellular immunogenicity. Furthermore, we identified the major negative regulator of IRF3-and NF-κB-dependent signaling, TNFAIP3 (A20), as a direct mRNA target of IGF2BP proteins and show that the immunosuppressive function of these RNA-binding proteins (RBPs) is, at least in part, mediated by their support of TNFAIP3 expression in myeloid leukemia cells.

## Results

### Downregulation of IGF2BPs expression activates ISRE-and NF-κB-dependent transcription with and without cytokine stimulation

Recently, we showed that genetic inhibition of IGF2BPs in mammalian cells leads to increase of ISGs expression, with and without cytokine stimulation. In this study, we investigated the role of these RNA-binding proteins in IRF3-and NF-κB-driven transcription using luciferase (Luc) and secreted alkaline phosphatase (SEAP) reporters. Without cytokine stimulation, genetic inhibition of IGF2BPs, either by shRNA knockdown (KD) or CRISPR/Cas9 knockout (KO) (Supplemental Fig. 1A, B), led to increased IRF3-driven expression of luciferase under the control of an interferon-sensitive response element (ISRE) promoter and NF-κB-driven alkaline phosphatase reporter (Fig. 1A). Type I interferon (IFN-β) treatment of cells with IGF2BP1, 2, and 3 KD yielded a significant increase of IRF3-dependent transcription compared to cytokine-stimulated non-targeting controls (Fig. 1A, data presented as Log2). Activation of IRF3 and NF-κB-induced signaling was accompanied by increased expression of ISGs (Fig. 1B) and increased phosphorylated IRF3 and NF-κB (Fig. 1C, D). The ISRE-driven reporter activation was partially rescued by co-transfection with siRNA against IFNAR1 (Fig. 1E), indicating IGF2BP-mediated effect on type I IFN signaling. Similarly, we observed an induction of ISRE-Luc and NF-κB-SEAP reporters using chemical inhibitors of IGF2BP1 (AVJ at [4-5μM]), and IGF2BP2 (IMP2-IN1 [20-25 μM] and CWI-1.2 [1-2 μM]) in HEK293T cells (Supplemental Fig. 1C, D). Therefore, we concluded that high levels of IGF2BP proteins suppress ISG expression by direct or indirect negative regulation of IRF3-and NF-κB-mediated transcription from ISRE and NF-κB-dependent promoters.

**Fig. 1.**
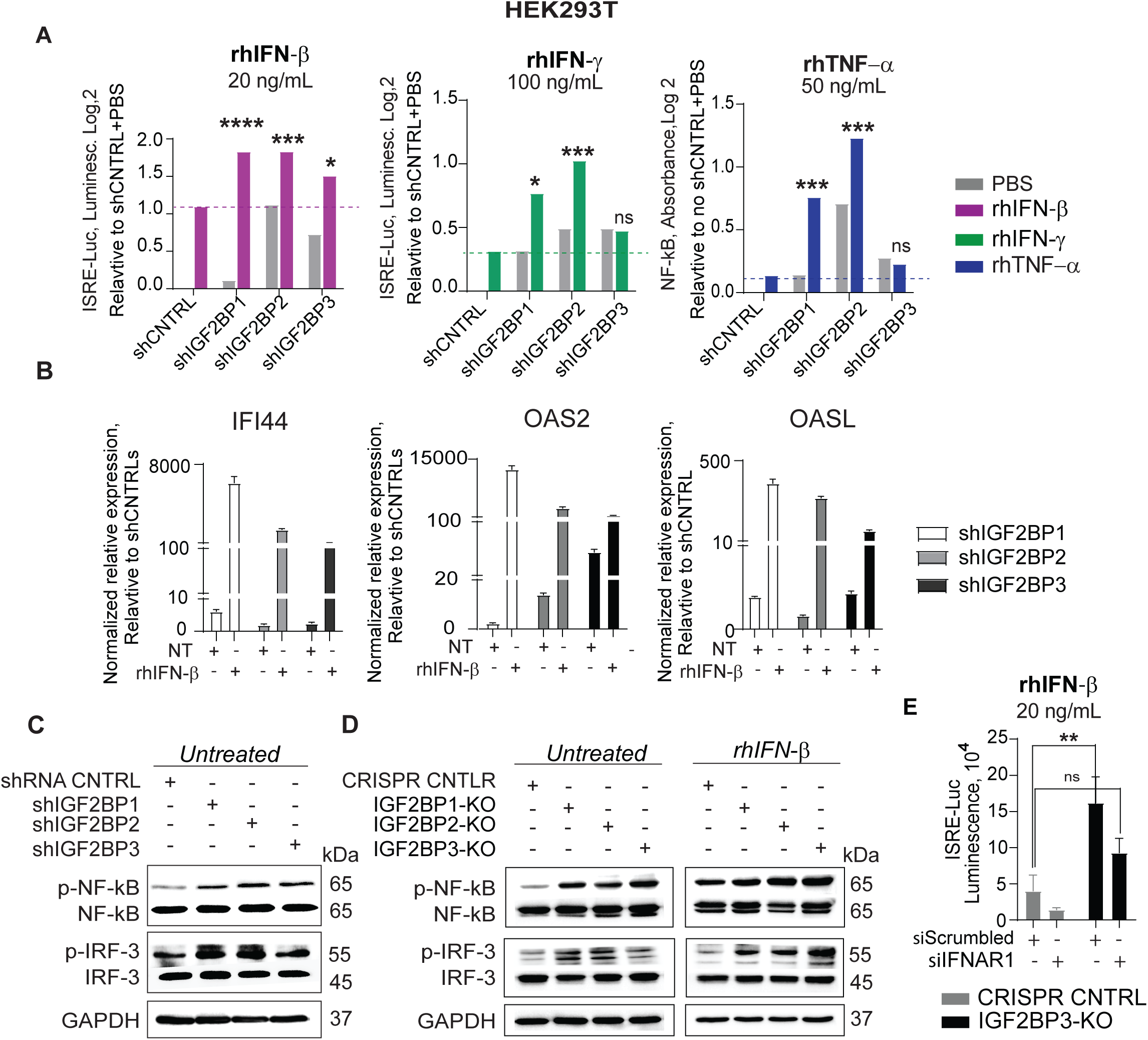
Downregulation of IGF2BPs expression activates ISRE-and NF-κB-dependent transcription with and without cytokine stimulation. **A** Representative analysis of ISRE-driven luciferase and NF-κB-driven secreted alkaline phosphatase (SEAP) activity in HEK293T cells with IGF2BPs-KD, relative to untreated/unstimulated non-targeting shRNA control (PBS), Log2 of original values (*n*=4). Indicated statistically significant difference between cytokine-treated IGF2BP-KD and cytokine-treated non-targeting control groups refers to original readouts and specified by the dashed line. **B** Representative analysis of interferon-stimulated genes’ expression by qRT-PCR in HEK293T cells non-treated (NT) or treated with rhIFN-β (20 ng/mL, 12 hours), transduced with shRNA against IGF2BP1-3. Data presented as normalized expression relative to corresponding non-targeting shRNA controls. Data is presented as mean ± SD (*n*=3). **C, D** Western blot analysis of endogenous levels of phosphorylated (p) and total IRF3 and NF-κB in IGF2BP1,2, and 3 gene knockdown (shRNA) and knockout (KO) clones, untreated or treated with rhIFN-β (20 ng/mL, 12 hours). **E** ISRE-driven luciferase in rhIFN-β-treated HEK293T cells with IGF2BP3-KO and non-targeting CRISPR/Cas9 control, co-transfected with siIFNAR1 or siScrambled RNA oligonucleotides. Data is presented as mean ± SD (*n*=3). Statistical analysis for all panels: unpaired *t*-test. ****p < 0.0001; \*\*\**p* < 0.001; \*\**p* < 0.01; \**p* < 0.05, not significant (ns).

### IGF2BP knockouts are associated with changes in chromatin accessibility and pan-transcriptome

Given previously and newly described role of IGF2BPs in regulating levels and activity of transcription factors, we hypothesized that these cytoplasmic RBPs may have indirect but significant effect on chromatin accessibility and transcription in the nucleus. To this end, we performed an integrative analysis with Assay for Transposase-Accessible Chromatin using sequencing (ATAC-seq) and RNA sequencing of subcellular fractions of HEK293T cells with IGF2BP1, 2, and 3-KO (Fig. 2A).

**Fig. 2.**
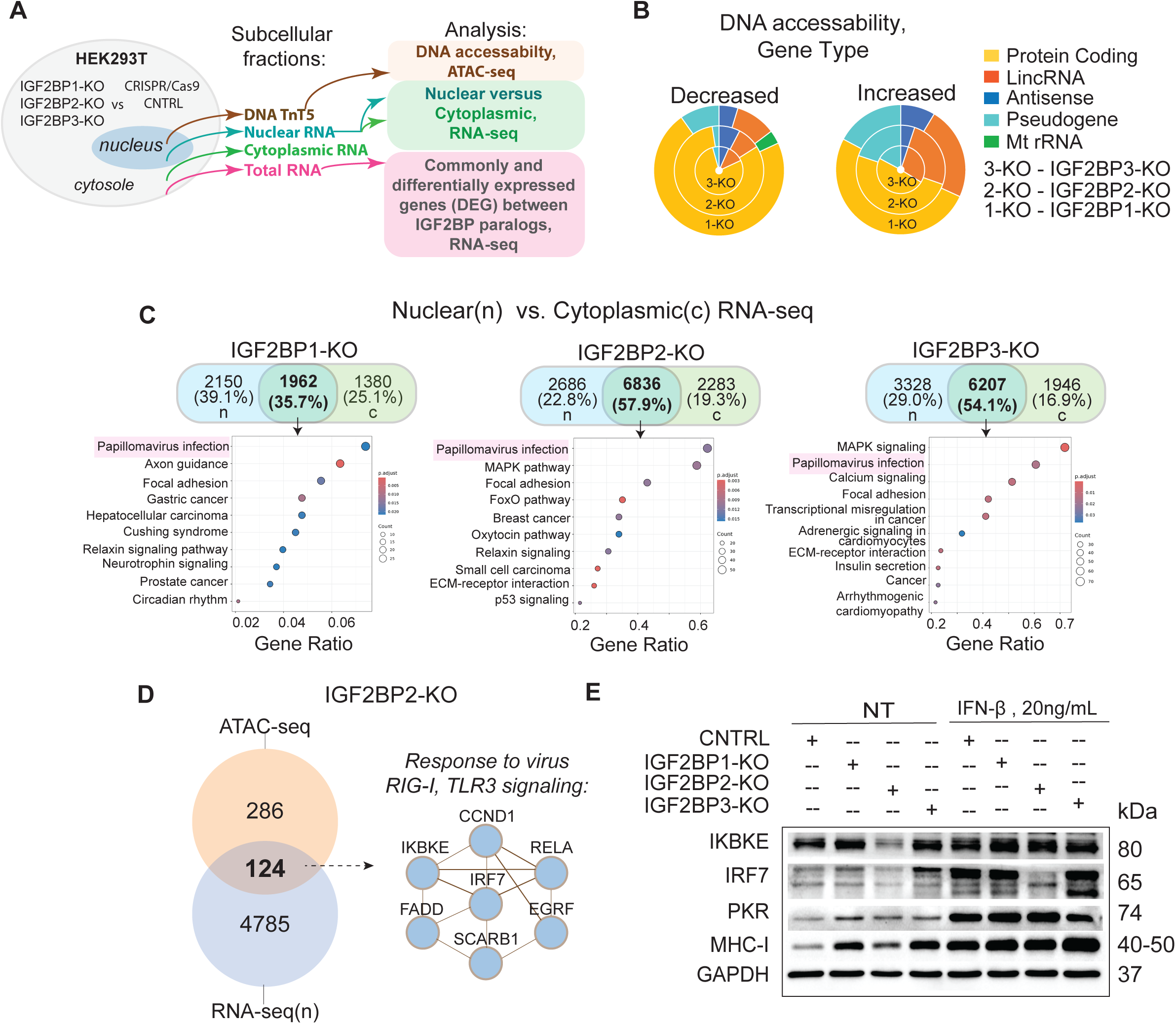
Integrative ATAC-seq and RNA-seq analysis of IGF2BPs knockout HEK293T cells. **A** Schematics of the experiment. **B** Gene types associated with decreased and increased chromatin accessibility in IGF2BP-KO cells. **C** VENN diagrams indicating common and unique DEGs in nuclear (n) and cytoplasmic (c) RNA fractions isolated from IGF2BP-KO. KEGG enrichment for upregulated DEGs shared between nuclear and cytoplasmic fractions for each IGF2BP-KO. **D** VENN diagrams indicating numbers of genes with significantly decreased chromatin accessibility measured by ATAC-seq analysis (n=286), and downregulated expression levels measured by RNA-seq analysis (n=4785) in IGF2BP2-KO HEK293T cells (left). Enrichment in protein domain structures in genes demonstrated both downregulated expression in RNA-seq and decreased DNA accessibility (*n*=124). **E** Protein levels of selected innate immunity regulators differentially expressed in IGF2BPs-KO; untreated (NT) or treated with rhIFN-β (20 ng/mL, 12 hours).

The ATAC-seq analysis identified genome loci with statistically significant changes in both decreased and increased DNA chromatin accessibility, representing ∼1-2% of the total genome. The numbers of chromatin areas with decreased accessibility were similar between IGF2BP-KOs: IGF2BP1-KO (566), IGF2BP2-KO (450) and IGF2BP3-KO (421) and mostly encompassed protein coding genes (Fig. 2B, Supplemental Fig. 2A, B, C). Regulators of endocytosis and protein transport were among common genes with decreased chromatin accessibility (Supplemental Fig. 3A). Increased chromatin accessibility data differ substantially between IGF2BP1-KO (172), IGF2BP2-KO (1,157) and IGF2BP3-KO cells (1,468) and were enriched with long intergenic non-coding RNA (lincRNA) and processed pseudogenes (Fig. 2B, Supplemental Fig. 2C). Genes encoding plasma membrane components were commonly affected by chromatin opening in IGF2BP-KOs (Supplemental Fig. 3C). Response to pathogens and immune cell interaction were among biological and molecular functions in ATAC-seq data for individual IGF2BP-KOs (Supplemental Fig. 3C, D).

Response to viral infection was one of the most upregulated pathways in total, nuclear, and shared nuclear and cytoplasmic fractions in all three IGF2BP-KOs (Fig. 2C, Supplemental Fig.4). The commonly upregulated genes included interferon-activated genes (e.g., *IFNAR1, ISG15, EIF2AK2 (PKR)*), transcriptional activators involved in inflammation (e.g., *CREB5*), and extracellular matrix proteins (e.g., *COL4A4, LAMA3, ITGA7, ITGB4*). Surprisingly, both ATAC-seq and RNA-seq identified downregulation of several key regulators of IRF-3-dependent transcription in IGF2BP2-KO (Fig. 2D, Supplemental Fig. 5). To validate the results of integrative ATAC-seq and RNA-seq analysis, we performed western blotting for several upregulated (e.g., PKR, MHC-I) or downregulated (e.g., IRF7, IKBKE) genes in IGF2BP-KO cells (Fig. 2E). Consistently, protein levels of both IRF7 and IKBKE were significantly decreased in IGF2BP2-KO cells. While expression of IKBKE was rescued by stimulation with IFN-β, expression of IRF7 was not restored with IFN-β treatment in IGF2BP2-KO cells. Protein levels of PKR and MHC class I were increased in all IGF2BP-KOs (Fig. 2E).

**Fig. 3.**
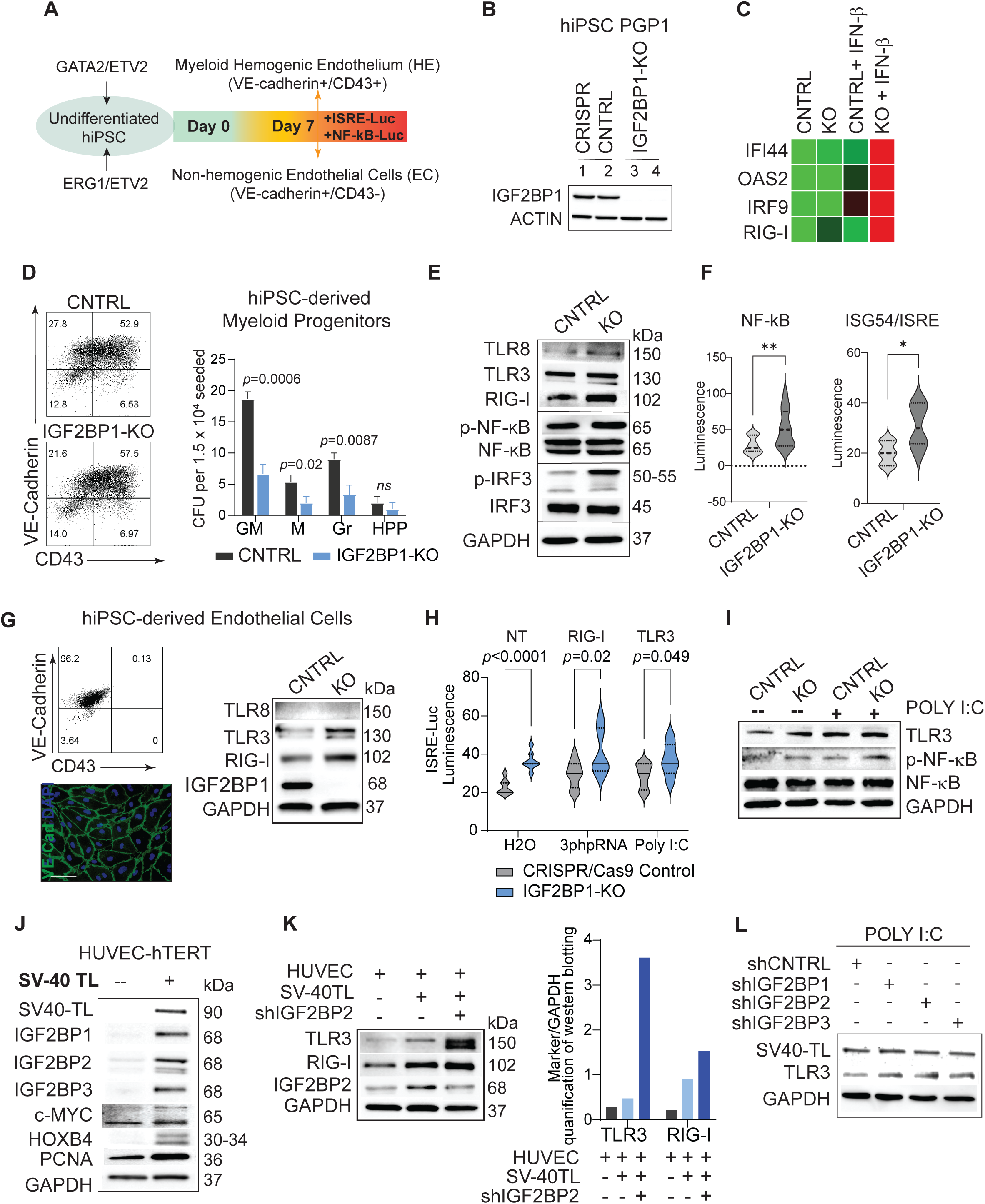
IGF2BPs suppress levels and activity of RNA sensors in embryonic-like and fetal hematoendothelial cells. **A** Schematic of the experiment. **B** Western blot analysis of IGF2BP1 expression in undifferentiated CRISPR/Cas9 control (CNTRL) and IGF2BP1-KO (KO) hiPSCs. **C** qRT-PCR analysis of endogenous ISGs expression in undifferentiated hiPSCs with and without rhIFN-β treatment (20 ng/mL, 12 hours). **D** Representative flow cytometric analysis of VE-cadherin and CD43 levels in hiPSC-derived myeloid HE (day 7, left), and corresponding colony forming units (CFU) assay, day 21: macrophage colonies (M), granulocyte (Gr), mixed granulocyte-macrophage (GM), high proliferative potential (HPP); data collected from three independent differentiation experiments. **E** Western blot analysis of endogenous expression of selected markers in myeloid cells, CRIPSR/Cas9 control versus IGF2BP1-KO, day 7. **F** Levels of intracellular NF-κB and ISRE-driven luciferase activity in myeloid cells, assessed by ONE-Glo^TM^ Luciferase assay (Promega), data are mean ± SD (*n*=3), indicated as the middle and side dashed lines. **G** Representative flow cytometric analysis and immunocytochemistry staining of VE-cadherin and CD43 levels non-hemogenic endothelial cells (ECs), (left); Western blot analysis of endogenous levels of TLR3, TLR8, and RIG-I expression in hiPSC-derived ECs (right). **H** ISRE-driven luciferase activity in EC differentiated from hiPSC non-targeting control and IGF2BP1-KO hiPSC, non-treated (NT) or treated with RIG-I and TLR3 agonist; Data are mean ± SD (*n*=3), indicated as the middle and side dashed lines. **I** Western blot analysis of TLR3 endogenous levels, phosphorylated and total NF-κB, and corresponding GAPDH in hiPSC-derived ECs treated with the TLR3 agonist poly(I:C) (10 μg/mL, 14 hrs). **J** Western blot analysis of IGF2BPs, c-MYC, HOXB4, PNCA, and SV40-T large antigen expression in HUVEC cells transfected with SV40-TL antigen expressing plasmid. **K** Western blot analysis and corresponding quantification of TLR3 and RIG-I levels in HUVEC cell lines transformed with SV40 T large antigen with and without IGF2BP2 knockdown. **L** Western blot analysis of TLR3 and SV40-T large antigen in HUVEC -SV40-TL expressing cells with IGF2BP1-3 knockdown and non-targeting control, treated with the TLR3 agonist poly(I:C), 10 μg/mL for 14 hrs. Statistical analysis for all panels: unpaired *t*-test. \*\**p* < 0.01; \**p* < 0.05, *n*=3.

**Fig. 4.**
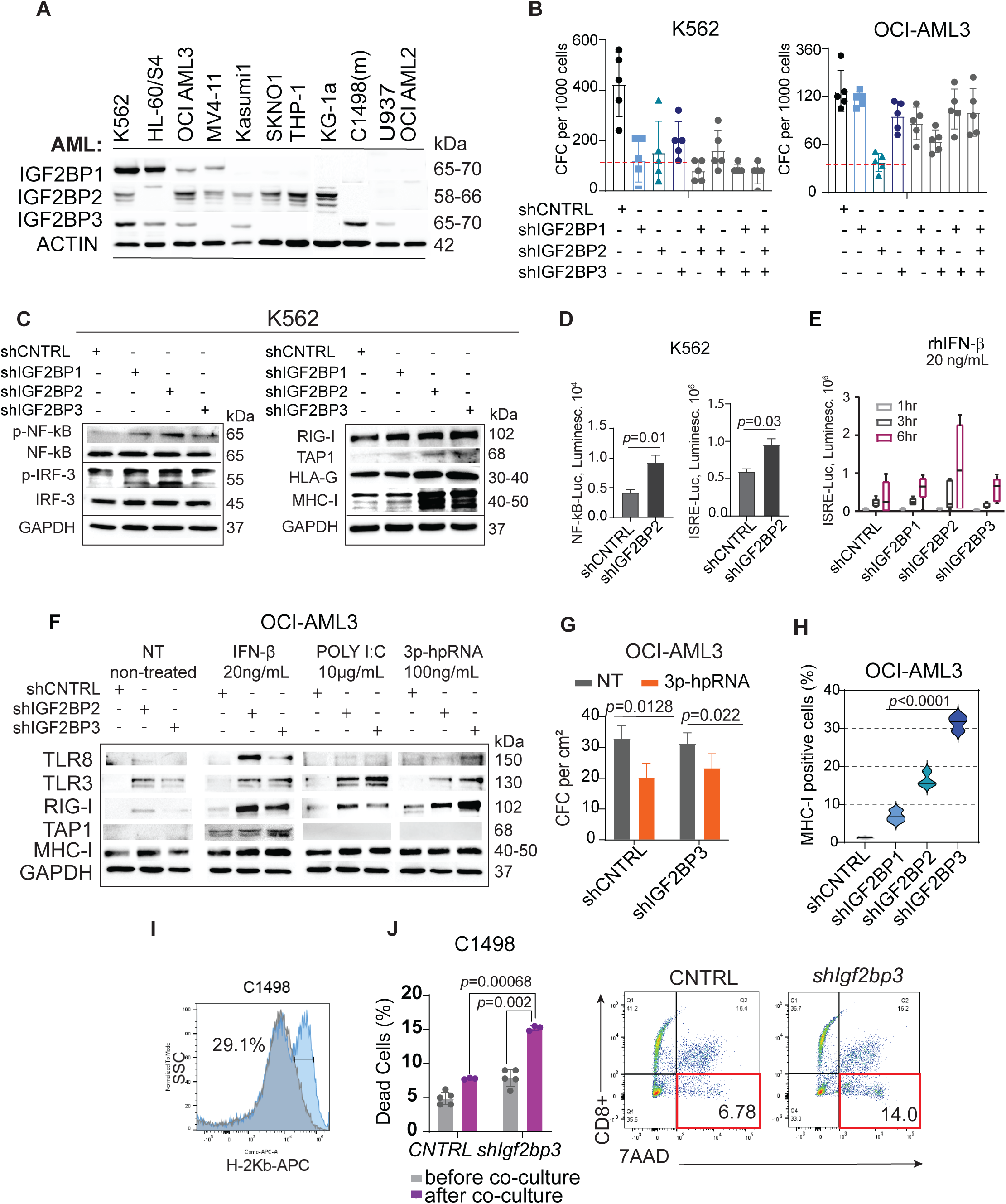
IGF2BPs expression in myeloid leukemia cells and their role in innate immune signaling. **A** IGF2BP1-3 protein levels in human AML cell lines and murine AML C1498. **B** Colony forming units (CFU) assay of K562 and OCI-AML3 cells transduced as indicated, day 21-post-transduction, Data is presented as mean ± SD (*n*=5). **C** Western blot analysis of endogenous protein levels in unstimulated K562 CML with IGF2BP1-3 KD. **D** Endogenous levels of IRF-3 and NF-kB-dependent transcription were assessed with Luciferase and SEAP reporters in untreated K562 cells with IGF2BP-2 KD. Data is presented as mean ± SD (n=3), statistical analysis was performed using two tailed t-test; **E** Time course assessment of intracellular expression levels of ISRE-driven luciferase in myeloid leukemia K562 cells with IGF2BPs-KD, treated with rhIFN-β (20 ng/mL, 1, 3, and 6 hours), Data is presented as mean ± SD (n=3). **F** Western blot analysis of OCI-AML3 with IGF2BP2-KD and IGF2BP3-KD, induced by IFN-β (20 ng/mL), TLR3 ligand poly(I:C) (10 μg/mL), RIG-I ligand 3p-hpRNA (100 ng/mL) for 12 hrs. **G** Colony forming units (CFU) assay for OCI-AML3 cells with shCNTRL and IGF2BP3-KD treated as indicated, day 21-post-transduction. Data is presented as mean ±SEM (*n*=3). **H** Quantification of cell surface MHC-I expression in OCI-AML3, assessed by flow cytometry, n=3. **I** Representative flow cytometric analysis of H-2K^b^ expression in murine AML C1498 with *Igf2bp3*-KD, day 7 post-selection with puromycin. **J** Quantification of 7AAD positive cells (%) and representative flow cytometric analysis of mouse C1498 AML with Igf2bp3-KD and non-targeting shRNA control, co-cultured with mouse T-cells, effector-to-target ratio 2:1, 48hrs (*n*=5). Statistical analysis between two samples assessed using two-tailed *t*-test or one-way ANOVA.

In summary, this analysis indicates that levels of cytoplasmic IGF2BP proteins exert a substantial impact on the pan-transcriptome with ∼2% overlap with changes in chromatin accessibility within or nearby differentially expressed genes. Our data suggest that *de novo* transcriptional activation and repression processes can occur because of depletion of IGF2BPs. Immunomodulatory functions of IGF2BP paralogs may vary depending on the cellular context or compensate for overactivation of pro-inflammatory signaling. Since nuclear transcriptomes of all three IGF2BP-KOs were enriched with genes regulating response to viral infection, we hypothesized that IGF2BPs attenuate the expression of ISGs by regulating the expression and/or activity of intracellular RNA sensors.

### IGF2BPs regulate endogenous PRR-mediated IRF3 and NF-κB signaling in embryonic-like and fetal hematoendothelial cells

Blood and endothelium are connective tissues with important immune and barrier functions in the body, expressing an array of pattern recognition receptors (Chen et al., 2025; Xu et al., 2022). To investigate the role of IGF2BPs in regulating the expression and activity of endogenously expressed single-stranded (ss) and double-stranded (ds) RNA sensors, such as TLR8, TLR3, and RIG-I, we utilized hiPSC-derived pan-myeloid hemogenic (VE-cadherin+/CD43+) and non-hemogenic (VE-cadherin+/CD43−) endothelial cells generated by forced expression of hematopoietic and endothelial transcription factors in IGF2BP1-knockout (KO) hiPSC clones (Fig. 3A, B). The stimulatory effect of IGF2BP1 knockout on type I IFN signaling was confirmed by qRT-PCR analysis of several interferon-stimulated genes (ISGs), including RIG-I, whose expression was markedly increased following IGF2BP1 depletion (Fig. 3C).

hiPSCs that lack IGF2BP1 expression produced a similar number of hematoendothelial VE-cadherin^+^/CD43^+^ progenitors compared to CRISPR/Cas9 control hiPSCs. However, their proliferative potential was significantly impaired (Fig. 3D). Western blot analysis indicated a substantial upregulation of RIG-I and p-IRF3 protein levels (Fig. 3E), accompanied by a significant increase of ISRE-and NF-kB-driven transcription in IGF2BP1-KO HE cells (Fig. 3F). IGF2BP1-KO endothelial cells expressed higher levels of TLR3 and RIG-I compared to CRISPR/Cas9 control (Fig. 3G). Similar to hiPSC-derived myeloid cells, endothelium lacking IGF2BP1 expression had a significant higher basal level of ISRE-driven reporter activity in unstimulated cells and increased sensitivity to RIG-I ligand 5’-triphosphate hairpin RNA (3p-hpRNA), and TLR3 ligand poly(I:C) (Fig. 3H, I).

Human umbilical vein primary endothelial cells (HUVEC) are of fetal origin. To maintain their *ex vivo* growth, HUVECs are immortalized by exogenous expression of human telomerase reverse transcriptase (hTERT) protein. HUVEC-hTERT cells express low levels of IGF2BP2 and almost undetectable protein levels of IGF2BP1 and IGF2BP3. We hypothesized that HUVEC-hTERT transformation by SV40-T large antigen (TL) will further increase HUVEC proliferation and induce expression of IGF2BPs. Indeed, SV40-TL transformation significantly increased or induced *de novo* endogenous expression of IGF2BP paralogs, proliferating cell nuclear antigen (PCNA), previously characterized IGF2BP1 and IGF2BP3 direct targets c-MYC and HOXB4 (Fig. 3J). SV40-TL-mediated transformation increased basal expression levels of TLR3 and RIG-I proteins which levels and activity were further enhanced by IGF2BP2 knockdown (Fig.3K, L).

In summary, these data indicate that endogenous expression of RIG-I, TLR3, and TLR8, as well as the ligand-induced activation of these receptors, negatively correlates with IGF2BP protein expression. Furthermore, mammalian cell transformation by the polyomavirus-derived SV40 large T antigen is associated with induction of the IGF2BP1–3 paralogs, which may contribute to suppression of innate immune responses by inhibiting the expression and activity of TLRs and RLRs.

To further test the hypothesis that high IGF2BP levels suppress activity of intracellular sensors, we utilized RLR and TLR expressing reporter systems in HEK293T cells co-expressing three IGF2BP paralogs. For initial screening, we examined endosomal ssRNA sensor TLR8, cytoplasmic dsRNA sensor RIG-I, and three non-RNA sensors: TLR5 which detects flagellin, NOD1 which detects the bacterial polypeptide L-Ala-γ-D-Glu-mDAP (Tri-DAP), and STING which detects the cyclic dinucleotide 2’3’-cGAMP. IGF2BPs expression was silenced using two sets of shRNAs targeting each IGF2BP paralog, and non-targeting shRNA controls.

A significant increase in ISRE-driven reporter activity was observed in the unstimulated RIG-I-expressing reporter cell line following genetic knockdown of each IGF2BP paralog (Supplemental Fig. 6A) and after treatment with the RIG-I agonist 3p-hpRNA (Supplemental Fig. 6B, C). The increased ISRE-regulated luciferase activity was completely rescued by co-treatment with the RIG-I antagonist and the TBK1 inhibitor Amlexanox in IGF2BP1-KD and IGF2BP2-KD cells and partially rescued in IGF2BP3-KD cells (Supplemental Fig. 6D). Similarly, unstimulated TLR8-expressing HEK293T cells with IGF2BP2 or IGF2BP3 knockdown exhibited significantly increased basal ISRE/ISG54-mediated luciferase activity (Supplemental Fig. 6E). A significant increase in NF-κB-driven reporter activity was also observed in IGF2BP3-KD cells treated with the ssRNA TLR8 agonist, and this effect was rescued by the TLR8-specific antagonist CU-CPT9a (Supplemental Fig. 6F). Finally, IGF2BP expression levels did not significantly affect STING-or NOD1-mediated signaling. Reduced NF-κB-driven reporter activity following treatment with the TLR5 agonist flagellin was observed in IGF2BP2-KD cells (Supplemental Fig. 7A), whereas no significant differences between the non-targeting control and IGF2BP1–3-KD cells were observed following STING or NOD1 stimulation (Supplemental Fig. 7B, C).

These data indicate that IGF2BP expression levels affect the activity of both cytoplasmic and endosomal RNA sensors. The IGF2BP family of proteins exerts a strong inhibitory effect on RIG-I signaling, both in the presence and absence of stimulation with its specific ligand. High levels of reporter protein expression positively correlate with endogenous levels of phosphorylated IRF3 and NF-κB, indicating activation of signaling pathways downstream of these receptors.

### Reactivation of embryonic IGF2BP1 and IGF2BP3 is essential for suppression of innate immune signaling pathways in AML

Previously, we showed that upregulation of IGF2BP1 and IGF2BP3 is associated with the leukemia stem cell phenotype defined by robust AML engraftment into NSG mice, high colony forming capacity (CFC), and high levels of ALDH1A1 expression (Elcheva et al., 2020b). Thus, K562 CML and HL-60/S4 AML, which express high levels of IGF2BP1 and IGF2BP3, had the highest leukemia stem cell score, whereas IGF2BP2 expressers (i.e., SKNO1 and Kasumi1) were among the least aggressive AML. Here, we further investigated the role of IGF2BP paralogs in the control of AML proliferation and innate immune signaling using a panel of AML cell lines and patient-derived AML (Figs. 4A, 5C).

**Fig. 5.**
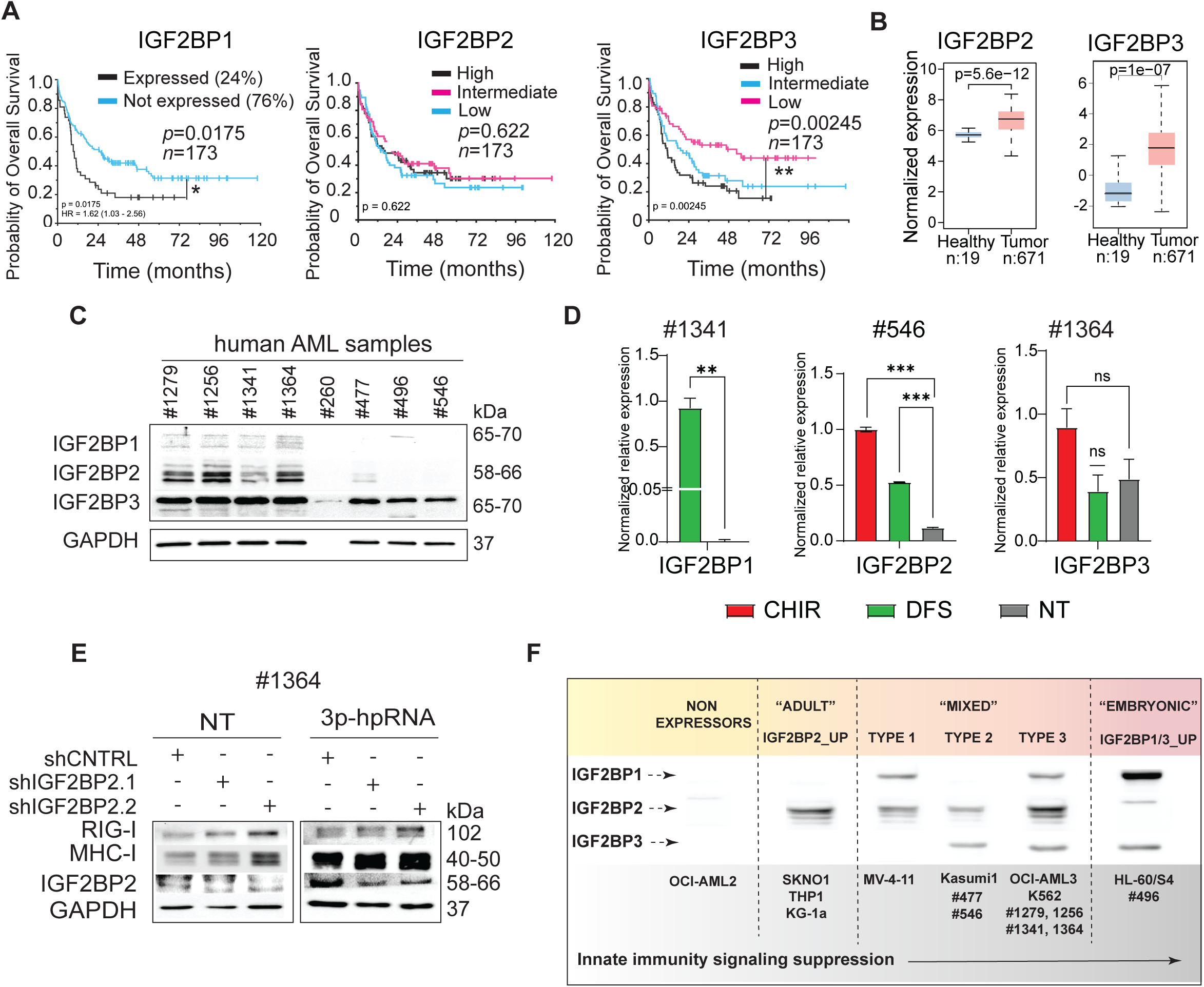
IGF2BP expression in primary AML and their role in innate immune signaling. **A** The overall survival (OS) of AML patients with various levels of IGF2BPs expression, TCGA database, assessed using log-rank tests. The statistically significant difference between AML expressing and not expressing IGF2BP1, and high and low levels of IGF2BP3 expression indicated as \*\**p* < 0.01; \**p* < 0.05. **B** Normalized IGF2BP2 gene expression (RPKM) in healthy individual bone marrow mononuclear cells compared with tumor samples obtained from patients (BeatAML-2.0 dataset). *P*-value is calculated using Welch’s unpaired two sample t-test, two sided, median values are indicated, box edges represent 25% and 75% quartiles. **C** Western blot analysis of IGF2BP proteins in primary AML samples. **D** Individual qPCR analysis of IGF2BPs levels in primary AML samples with and without disulfiram (DSF) (1μM for 24hrs) or CHIR99021 treatment (10μM for 48hrs). **E** Western blot analysis of RIG-I and MHC-I in primary AML cells with IGF2BP2-KD (shRNA sequence 1 and 2) treated as indicated. **F** Proposed role of IGF2BPs expression in AML and innate immune signaling.

**Fig. 6.**
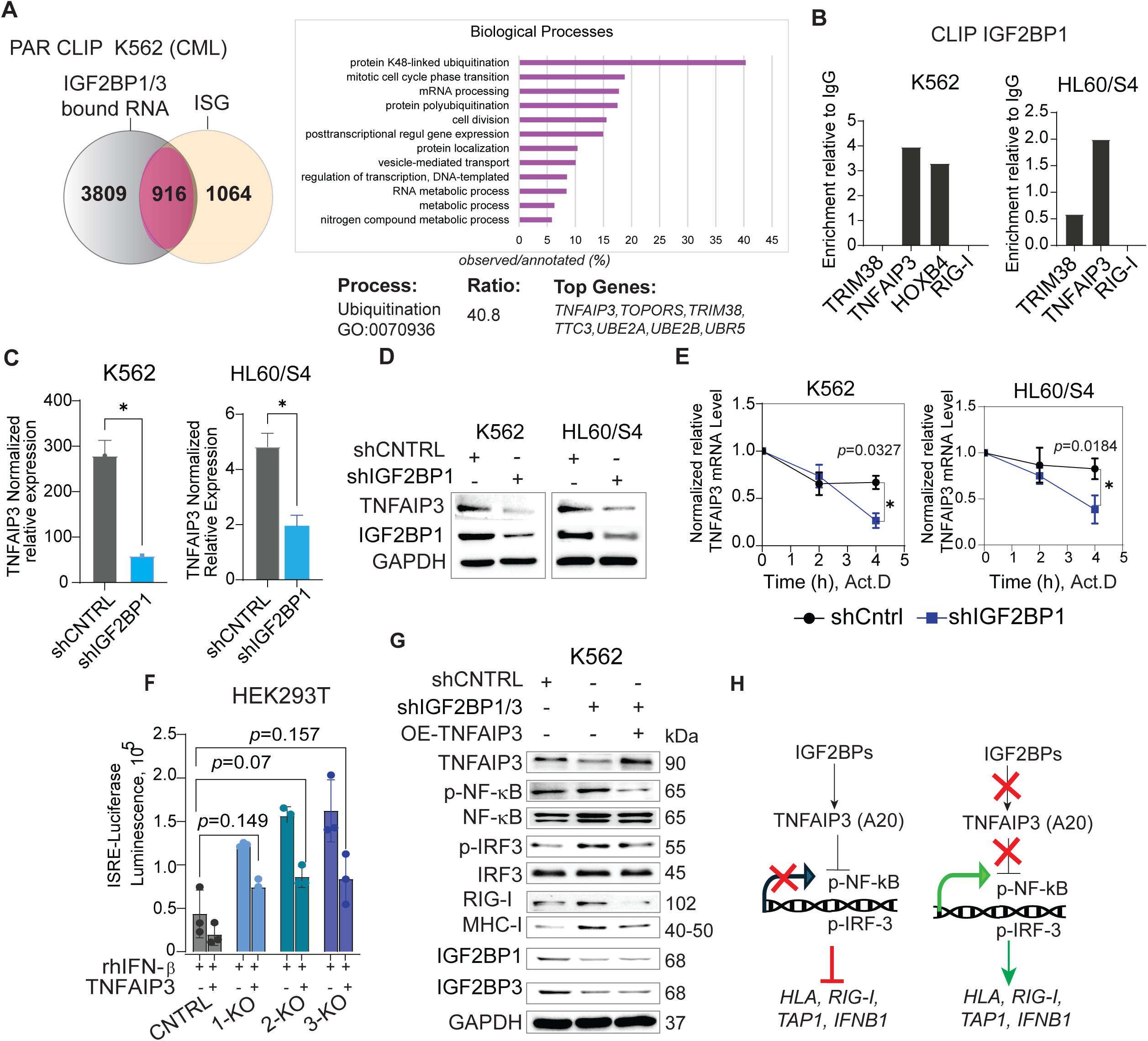
IGF2BPs suppress innate immune signaling by supporting TNFAIP3 expression. **A** VENN diagram for datasets generated from photoactivatable-ribonucleoside-enhanced crosslinking and immunoprecipitation (PAR CLIP) in K562 CML and ISGs annotated in GeneCards database (left). GO pathway analysis of direct mRNA IGF2BP1 targets in PAR CLIP (n=916) with highest ratio for genes observed to genes annotated belongs to K-48 ubiquitination (40.8%, GO:0070936), the top observed genes are as indicated. **B** qRT-PCR analysis of select genes analyzed in K562 and HL60/S4 AML by RNA-protein crosslinking and immunoprecipitation (CLIP) assay, enriched in IGF2BP1 precipitated versus IgG immunoprecipitation control. **C** qRT-PCR analysis of TNFAIP3 total mRNA levels in K562 and HL60/S4 AML with IGF2BP1-KD. **D** Western blot analysis of TNFAIP3 total protein levels of K562 and HL60/S4 AML with IGF2BP1-KD. **E** qRT-PCR analysis of TNFAIP3 mRNA levels in K562 and HL60/S4 AML with IGF2BP1-KD or non-targeting RNA control, treated with Actinomycin D (1μg/mL) for indicated number of hours. **F** ISRE-luciferase assay in HEK293T cells treated with rhIFN-β, co-expressing non-targeting shRNA control or shRNA against IGF2BP1, 2, and 3 in combination with TNFAIP3 overexpression. Vehicle control (VC) cells contain corresponding negative controls (PBS, siControl, or empty vector). Data are mean ± SD (*n*=3). **G** Overexpression (OE) of TNFAIP3 rescues effect of IGF2BP1 and IGF2BP3 double knockdown in K562 CML, assessed by western blotting. **H** Proposed mechanism of IGF2BPs/TNFAIP3 suppression of innate immune signaling.

First, we assessed whether any of the co-expressed IGF2BP paralogs played a dominant role in maintaining AML growth and proliferation. To this end, we performed single and multiple IGF2BP gene knockdowns in K562 and OCI-AML3 cells (Fig. 4B). While some additive effect was observed in K562 cells, no additive or synergistic effect occurred in double and triple gene knockdowns in OCI-AML3 cells, suggesting that the proliferative potential of AML is dependent on one of the most abundantly expressed paralogs. However, the immunomodulatory functions of IGF2BPs may not coincide with their pro-proliferative role. While K562 proliferation depends mostly on IGF2BP1 and less on IGF2BP2 and IGF2BP3, endogenous priming of IRF3 and NF-κB-driven transcription was evident in all IGF2BP-KD cells (Fig. 4C, D). Downregulation of IGF2BP2, but not IGF2BP1, yielded the strongest induction of ISRE-luciferase expression upon rhIFN-β treatment (Fig. 4E). Similarly, OCI-AML3 treatment with rhIFN-β, the TLR3 ligand poly(I:C), or the RIG-I ligand 3p-hpRNA, led to a robust increase of TAP1, MHC-I, TLR3, TLR8, and RIG-I levels in both IGF2BP2-KD and IGF2BP3-KD cells (Fig. 4F), but did not increase the anti-proliferative effect of IGF2BP3 gene knockdown upon RIG-I induction (Fig. 4G). Therefore, we conclude that the pro-proliferative and anti-inflammatory functions of IGF2BPs might be conveyed by different paralogs simultaneously co-expressed in a cell. Genetic inhibition of IGF2BP3 in human and mouse AML increased cell surface levels of MHC-I up to 30% (Fig.4H, I), and significantly improved AML killing in mouse syngeneic co-cultures with HLA-matching T-cells (Fig.4J).

We then examined SKNO1, THP-1, and KG-1a AML cell lines that express only or mostly IGF2BP2, the expression pattern associated with normal adult hematopoiesis. In these cells, IGF2BP2 genetic knockdown led to either none or partial activation of innate immune signaling, often accompanied by massive cell death (Supplemental Fig.9A-D). Limited responses of IGF2BP2 expressers suggest that reactivation of embryogenesis-specific IGF2BP1 and IGF2BP3 maximize AML resistance to PRR agonists and are essential for the suppression of innate immune signaling pathways in AML. We noted that IGF2BP2 protein expression is increased in response to IFN-β, RLR, and TLR agonists, suggesting that IGF2BP2 and its paralogs belong to ISG (Supplemental Fig. 8 D, E, F).

The analysis of IGF2BP1-3 levels in normal hematopoietic stem cells (HSCs) indicate that IGF2BP2 is expressed in normal CD34^+^HSCs, while IGF2BP3 levels are very low and lGF2BP1 is almost undetectable (Supplemental Fig. 9A)(Glaß and Hüttelmaier, 2023) Respectively, high levels of IGF2BP2 and IGF2BP3 were detected in all AML subjects, whereas IGF2BP1 expression was present in 24% of myeloid leukemias (Fig. 5A). Upregulation of IGF2BP1 and IGF2BP3 was associated with worse survival. Patients with intermediate and low levels of IGF2BP2 expression exhibit higher life expectancy than patients with IGF2BP2 upregulation, but no statistical significance was determined in the TCGA data set (central graph). The analysis of a larger AML cohort (TARGET) showed a significant association of a shorter overall survival (OS) in patients with high levels of IGF2BP2 (data not shown), which was also significantly upregulated in myeloblasts compared to normal bone marrow progenitors (Fig. 5B). In summary, these data indicate that upregulation of IGF2BP expression in AML, especially oncofetal IGF2BP1 and IGF2BP3 paralogs, is associated with unfavorable prognosis and worse clinical outcomes. Interestingly, IGF2BP3, but not IGF2BP2, was the most abundantly expressed paralog in primary human AML samples (Fig. 5C), supporting observation that IGF2BP3 upregulation represents a leukemogenic mechanism in the majority of AML cases (Zhang et al., 2022).

Given the established roles of WNT/β-catenin and hypoxia signaling in maintaining the leukemia stem cell phenotype and activating IGF2BPs, we examined IGF2BP expression levels in patient-derived AML samples before and after treatment with the GSK-3β inhibitor CHIR99021 and disulfiram (DSF), which activate β-catenin and hypoxia-inducible factor 1α (HIF-1α) signaling, respectively (Deynoux et al., 2016; Pepe et al., 2022). Indeed, IGF2BP1-3 expression significantly increased in the majority of primary AML samples (Fig. 5D, Supplemental Fig. 9 B, C). Genetic inhibition of IGF2BPs in primary AML led to upregulation of MHC-I and RIG-I protein expression, with and without agonist treatment, similar to AML cell lines co-expressing IGF2BP1-3 (Fig.5E). These data indicate that suppression of innate immune signaling is maximized by the co-expression of IGF2BP paralogs, which may promote leukemia development through immune evasion (Fig. 5F).

### IGF2BPs suppress innate immune signaling by promoting TNFAIP3 expression

Taking into consideration mRNA-binding properties of IGF2BPs, the increase of ISGs expression, including RIG-I, TLR3, TLR8 and MHC-I, in IGF2BP loss-of-function systems raises a question about the molecular mechanisms of their antagonistic expression patterns, in particular, if there is a direct binding of IGF2BPs with ISG mRNAs. To address this question, we aligned data from previously performed RNA-protein crosslink and immunoprecipitation in K562 myeloid leukemia cells with more than 2000 interferon-stimulated protein coding genes annotated in GeneCards database (Fig. 5A). The Gene Ontology (GO) analysis for biological process enrichment revealed that roughly half of ISG mRNA transcripts (n=916) showed binding with IGF2BPs regulating RNA and protein expression (e.g., protein K48-linked ubiquitination), cell cycle, vesicle-mediated transport, and transcription (Fig.5A). Genes upregulated in response to virus infection, innate immune and cytokine responses (e.g., OAS1, OAS2, IFI6, IFI16, ISG2, ISG15, IFIT1-3, 5 etc.,) are not likely to be regulated by binding with IGF2BPs (n=1064, VENN diagram Fig. 5A). Notably, cytosolic PRR mRNAs such as RIG-I (DDX58), TLR3, TLR8 showed no binding with IGF2BPs in K562 PAR CLIP data and several RNA-protein binding datasets (Supplemental Fig. 10).

To confirm these findings, we performed CLIP analysis to assess IGF2BP1 binding to the mRNAs of the previously characterized IGF2BP1 target HOXB4, RIG-I, and the two negative regulators of RIG-I signaling, TNFAIP3 and TRIM38, whose mRNAs contain IGF2BP1–3 binding sites identified by PAR-CLIP. Both K562 and HL60/S4 ribonucleoprotein immunoprecipitates were negative for RIG-I mRNA but were enriched for TNFAIP3 and HOXB4 transcripts (Fig. 5B). Total TNFAIP3 mRNA and protein levels were reduced in IGF2BP1-KD K562 and HL60/S4 cells (Fig. 5C, D). An actinomycin D chase assay showed a significant decrease in TNFAIP3 mRNA stability in IGF2BP1-depleted leukemia cells, indicating that IGF2BP1 directly binds to and stabilizes TNFAIP3 mRNA (Fig. 5E). TNFAIP3 overexpression in IGF2BP1–3-KO clones at least partially rescued ISRE-luciferase reporter activity (Fig. 5F). Likewise, the IGF2BP1/3-KD-mediated increases in phosphorylated IRF3 (p-IRF3), phosphorylated NF-κB (p-NF-κB), RIG-I, and MHC class I protein levels were partially rescued by TNFAIP3 overexpression in K562 chronic myeloid leukemia (CML) cells (Fig. 5G). In summary, we identified TNFAIP3, a major negative regulator of IRF3-and NF-κB-dependent signaling, as a direct mRNA target of the IGF2BP proteins. Based on these findings, we propose a negative-feedback mechanism in which IGF2BPs suppress innate immune signaling by positively regulating TNFAIP3 mRNA stability and protein expression in response to activation of interferon signaling (Fig. 5J).

## Discussion

The IGF2BPs were discovered in the context of embryonic development and cancer progression as oncofetal proteins expressed in rapidly developing, growing tissues (Bell et al., 2013; Degrauwe et al., 2016; Hansen et al., 2004). Upregulation of IGF2BPs was detected in various types of cancer and was associated with mRNA stability and translation of key transcriptional regulators of proliferation and self-renewal (e.g., *c-MYC*, *KRAS, HOXB4*, *MYB*) (Doyle et al., 1998; Elcheva et al., 2020a; Mackedenski et al., 2018). Recently, we discovered a dual function of IGF2BP proteins that, in addition to promoting proliferation, suppress expression of interferon stimulated genes in normal and transformed mammalian cells (Elcheva et al., 2023).

In this study, we examined the underlying mechanisms of IGF2BP-dependent regulation of innate immune responses in phenotypically normal, transformed, and cancerous cells co-expressing embryonic IGF2BP1-3 paralogs. We showed that IGF2BPs expression is negatively correlated with IRF3-and NF-κB-mediated transcription from ISRE-and NF-κB-dependent reporters. Importantly, IGF2BPs depletion leads to endogenous activation of IRF3-and NF-κB-controlled transcription in the absence of intrinsic and extrinsic molecular triggers such as stimulation with cytokines or PRR agonists. The enrichment of nuclear cell compartments with mRNA transcripts associated with viral infection, PI3K-Akt, and MAPK pathways in IGF2BP-KOs suggests *de novo* transcription in response to deficiency of these proteins. Our integrative analysis of chromatin accessibility and gene expression showed that significant portions of the transcriptome are uniquely regulated by individual IGF2BP paralogs, suggesting that IGF2BP family members may function complementarily to fine-tune innate immune responses and prevent excessive pro-inflammatory signaling. Moreover, pro-proliferative and immune-modulatory functions of IGF2BPs might be conveyed by different paralogs simultaneously co-expressed in a cell.

Strong association of IGF2BP deficiency with anti-viral signaling activation prompted us to investigate the role of IGF2BPs in the levels and activity of cytoplasmic PRRs. Genetic inhibition of IGF2BPs was accompanied by upregulation of RIG-I, TLR3, and TLR8 protein levels, increased sensitivity to their specific agonists and robust activation of downstream IRF-3 and/or NF-kB transcription. This phenotype was particularly evident in embryonic-like hematoendothelial cells and human AML cells co-expressing IGF2BP1 and IGF2BP3, suggesting that the embryonic-like transcriptome, of which IGF2BP1 and IGF2BP3 are integral components, is associated with reduced intracellular immunogenicity. Indeed, several studies show that programs of pluripotency and interferon signaling are incompatible (Eggenberger et al., 2019), (Wu et al., 2018). In support of these observations, our bioinformatics analysis of three IGF2BP-RNA-bindning data sets: eCLIP for IGF2BP1, 2 and 3 in hiPSCs (Conway et al., 2016), PAR CLIP in HEK293T (Hafner et al., 2010), and PAR CLIP in K562 (Elcheva et al., 2020a), indicate that hundreds of messengers encoding cytokines, chemokines, and transcriptional regulators of anti-viral responses have no binding with IGF2BP family of proteins. Whereas mRNAs of interferon-stimulated genes encoding post-transcriptional regulators of gene expression, protein polyubiquitination, mRNA processing, DNA-templated transcription, and cell division are direct targets of IGF2BP paralogs. Specifically, protein K48-linked ubiquitination was among most enriched biological processes among 916 IGF2BP-bound interferon-responsive mRNA targets in K562. Our CLIP assay of K562 and HL60/S4 myeloid leukemia cells confirmed direct binding with one of the enriched targets, TNFAIP3 (A20).

TNFAIP3 is a major negative regulator of both IRF-3 and NF-kB driven transcription via K48-linked and K63-linked ubiquitination (Parvatiyar et al., 2010) (Catrysse et al., 2014). Our study shows that RNA and protein levels of TNFAIP3 are positively regulated by IGF2BP paralogs. Increase of IRF-3 and NF-kB-dependent transcription and expression of RIG-I and MHC-I, upregulated in response to IGF2BP1-3 genetic inhibition, were rescued by forced expression of TNFAIP3. In addition, we showed that transcriptional activation of ISRE and NF-kB-controlled reporters, in response to IGF2BP genetic inhibition, can be efficiently rescued by a PRR-specific antagonist (e.g., TLR8 blocker CU-CPT9a) and the inhibitor of the downstream TANK-binding kinase1 (TBK1) and nuclear factor kB kinase epsilon (IKKε) (e.g., Amlexanox), which supports our hypothesis that IGF2BP family of proteins suppress activity of cytosolic RNA sensors and downstream IRF-3 and NF-kB-dependent transcription.

Expression of TNFAIP3 is shown to be activated by NF-κB which provides negative feedback regulation to damp inflammation (Vereecke et al., 2009). Interestingly, IGF2BPs are listed as interferon stimulated genes in the GeneCard database as well as TNFAIP3. In our study, IGF2BP2 protein levels increase with interferon treatment in AML, suggesting that IGF2BP2 and its paralogs are ISGs. In this case, the increased levels of IGF2BPs in response to interferon and PRR stimuli would initiate negative regulatory feedback loop, reducing innate immune signaling, and maintaining cell cycle progression by supporting TNFAIP3 expression. IGF2BP-mediated suppression of PRRs levels and activity would also be beneficial for viral transformation as we show by SV-40TL expression. The observation that IGF2BPs “fuel viral carcinogenesis” (Glaß and Hüttelmaier, 2023) is consistent with our study.

Our findings provide new perspective and potential new strategies for the treatment of leukemia and other types of cancers with high levels of IGF2BP expression. RIG-I and TLR3 agonists are being actively investigated as anti-cancer therapeutic agents (Elion and Cook, 2018; Yang et al., 2022). It has been shown that activation of RIG-I is strongly associated with HLA expression and MHC class I antigen presentation (Such et al., 2020; Thier and Paschen, 2021) and increases AML responsiveness to inhibitors of checkpoint blockade in an animal model (Ruzicka et al., 2020). Given that MHC-I is downregulated in 40-90% of human cancers (Cornel et al., 2020), IGF2BPs treatments could enhance antigen presentation and increase patient responsiveness to immunogenic therapies (Cornel et al., 2020). Not limited to carcinogenesis, understanding the role of IGF2BPs in the modulation of the levels and activity of PRRs may shed new light on the development of autoimmune and neurodegenerative diseases with aberrant innate immune signaling.

### Data and code availability

Data used for bioinformatics analysis were deposited in NCBI Gene Expression Omnibus (GEO) repository and can be accessed through the series accession numbers: GSE311641 and GSE311644 (ATAC-seq and RNA-seq integrative analysis, HEK293T), GSE138063 (PAR CLIP K562), GSE21918 (PAR CLIP HEK293), GSE78509 (eCLIP hPSCs).

## Acknowledgments

We are thankful to Dr. Sirisha Pochareddy and Genome Sciences Core’s personnel for assistance. Genome Sciences Core (RRID:SCR_021123) services and instruments used in this project were funded, in part, by the Pennsylvania State University College of Medicine via the Office of the Vice Dean of Research and Graduate Students and the Pennsylvania Department of Health using Tobacco Settlement Funds (CURE). We are also grateful for assistance provided by Genomics Research Incubator (RRID:SCR_024530), a collaborative research space and an initiative of the Eberly College of Science and the Huck Institutes of Life Sciences at the Pennsylvania State University. The content is solely the responsibility of the authors and does not necessarily represent the official views of the University or College of Medicine. The Pennsylvania Department of Health specifically disclaims responsibility for any analyses, interpretations or conclusions.

## Author contributions

IE developed a concept, designed and performed experiments, analyzed data, made figures and wrote the paper; GK, RA performed and analyzed experiments, contributed to figures and manuscript preparation; ZL, AM, KK, YU, SM performed bioinformatics analysis and made figures; AP, MH performed and analyzed PAR-clip data; MW, SK, CK, TS, JH, GLM, JZ, AS, HZ, EH assisted with experimental design, procedures, data analysis, and manuscript preparation; all authors contributed to manuscript editing.

---

**Supplemental Fig. 1.**
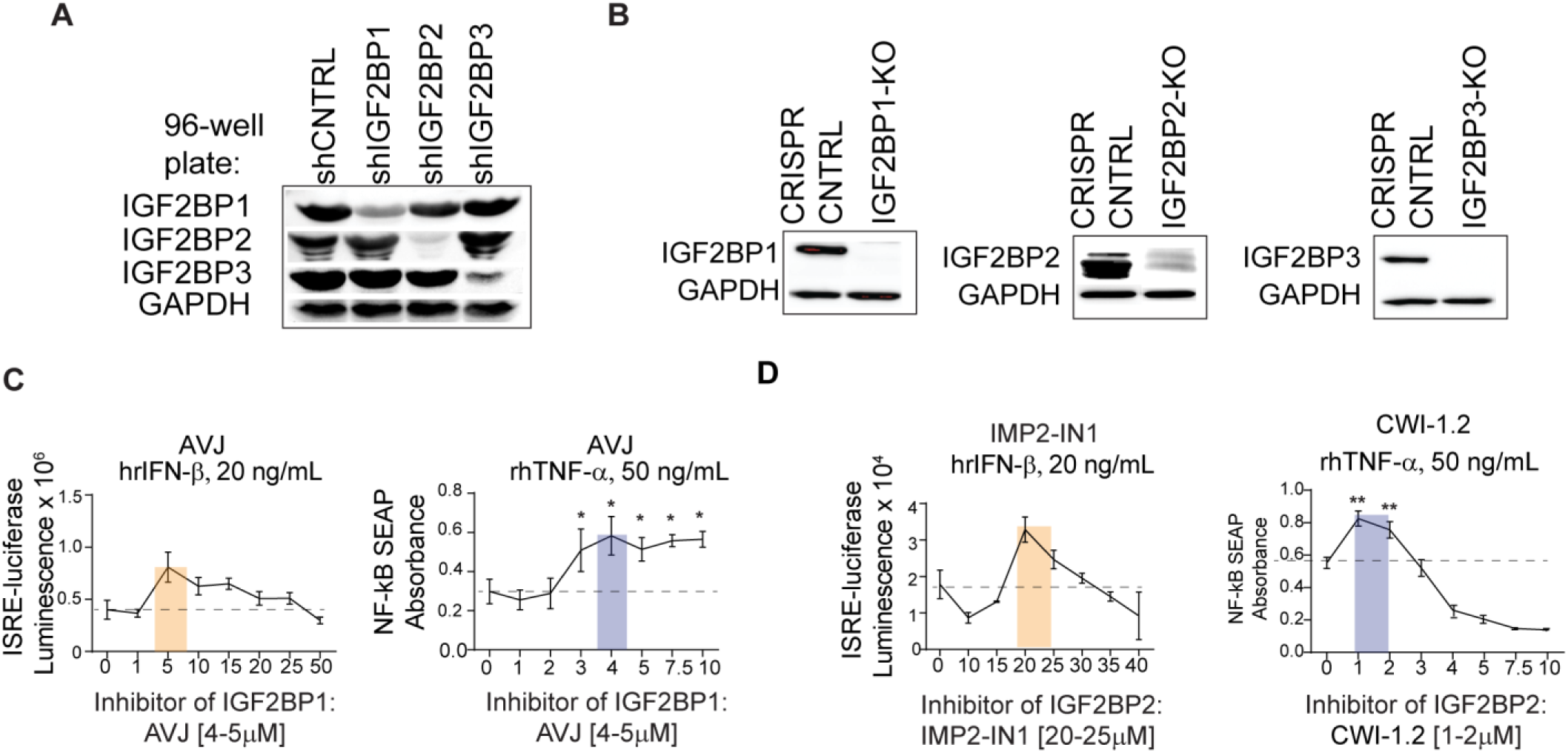
(Supplemental to main Fig.1): Inhibition of IGF2BPs expression activates ISRE-and NF-kB-dependent transcription. **A** Western blot analysis of IGF2BP levels in HEK293T cells expressing secreted Luc and SEAP reporters. HEK293T cells were stably expressing non-targeting shRNA controls and shIGF2BP1, 2, or 3 targeting shRNA (IGF2BP gene knockdown, KD). Cells were seeded for the plate reader assays and proteins were isolated from 3 wells of 96-well plate right after reading of luminescence/absorbance; **B** Western blot analysis of IGF2BP levels in IGF2BP gene knockout (KO), single cell-derived clones of HEK293T cells, transfected with CRISPR/Cas9-GFP non-targeting control or CRISPR/Cas9-GFP gRNA against IGF2BP1-3; **C, D** Luciferase and SEAP activity in HEK293T dual cells expressing both reporters, treated with indicated doses of IGF2BP1 and IGF2BP2 inhibitors. Data are presented as mean ± SD (*n*=3), two-tailed *t*-test \*\**p* < 0.01; \**p* < 0.05 between the basal activity of untreated (0μM) samples and treated cells, marked with a threshold. Numbers in brackets indicate concentrations inducing reporter activity, which are also highlighted within plots.

**Supplemental Fig. 2.**
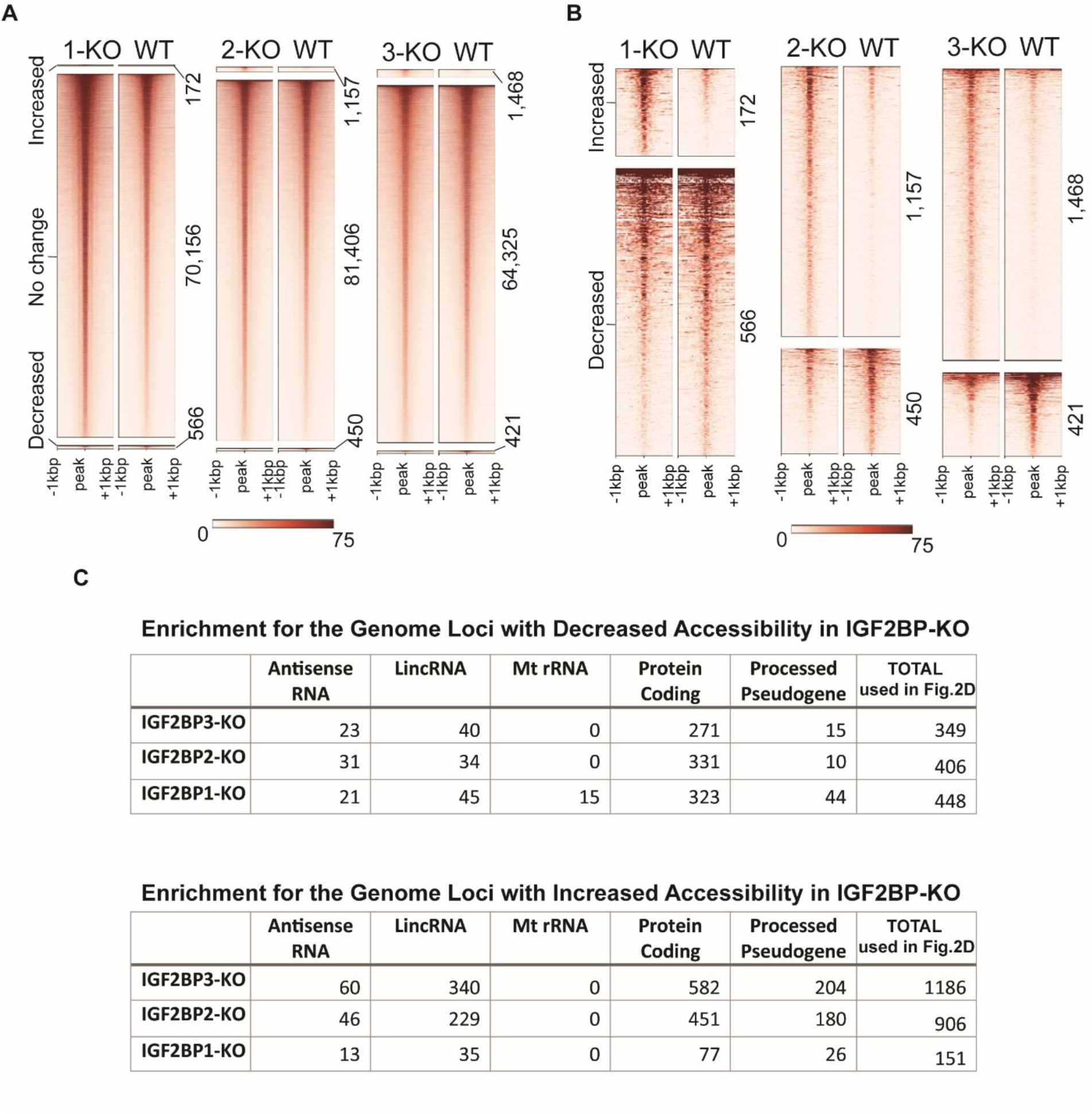
(Supplemental to main Fig.2). ATAC-seq analysis of IGF2BP-KO in HEK293T cells. **A** Mapping of chromatin accessibility by ATAC-seq in IGF2BP1-KO (1-KO), IGF2BP2-KO (2-KO), and IGF2BP3-KO (3-KO) cells compared to CRISPR/Cas9 non-targeting control (“wild type”, WT): total number of identified chromatin loci with significant increased, decreased, or no significant change (no change) in DNA accessibility. **B** Heatmap visualization of ATAC-seq profiling of statistically significant increased and decreased chromatin accessibility in IGF2BP-KO cells also shown in 2B as increased and decreased loci. **C** Types of genome loci with decreased or increased accessibility that are most frequently represented in ATAC-seq analysis of IGF2BP-KO.

**Supplemental Fig. 3.**
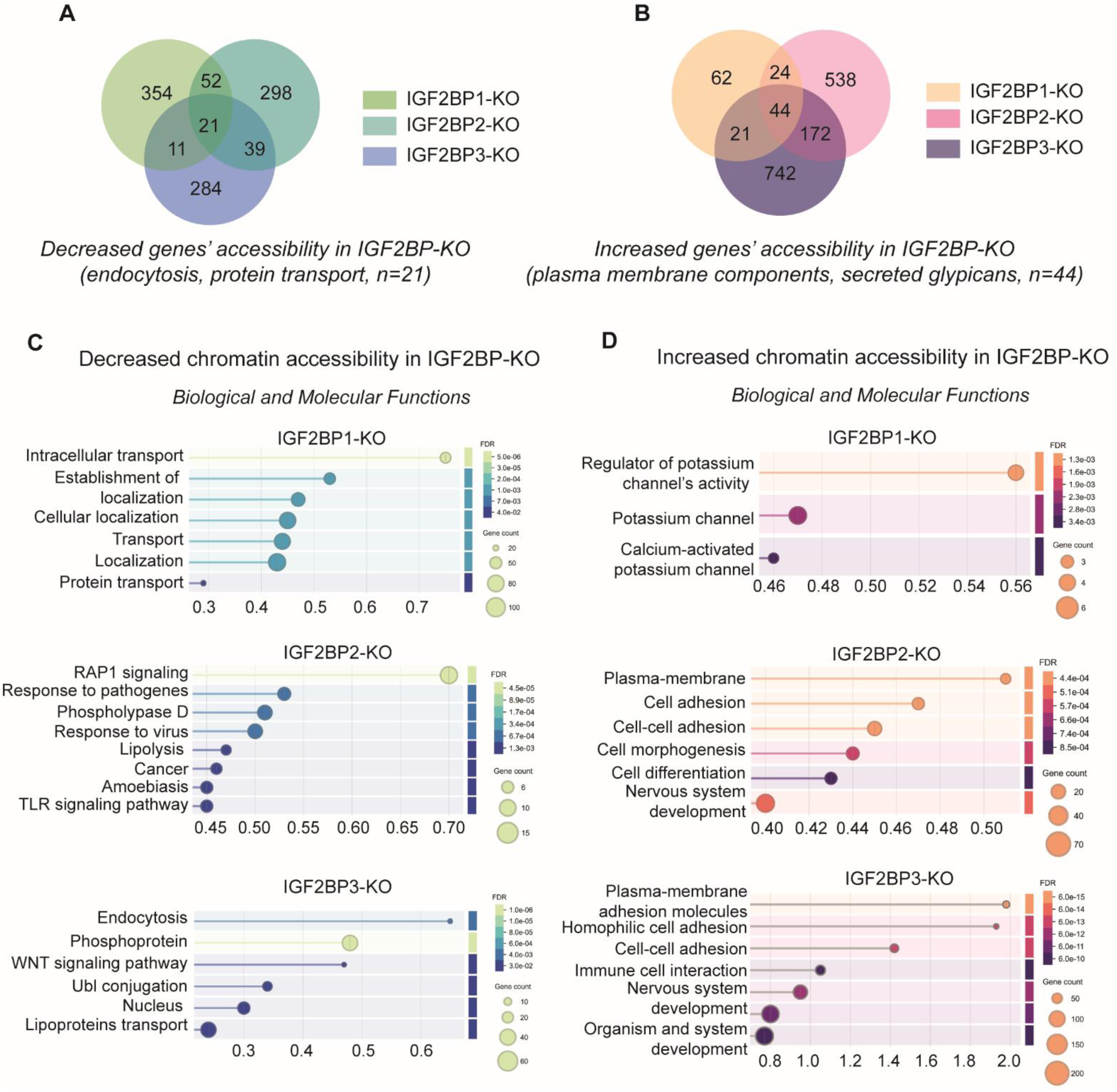
(Supplemental to Main Fig.2). Gene ontology analysis for genes with increased or decreased chromatin accessibility in IGF2BP-KO. **A** VENN diagram for genes associated with the significant decrease of chromatin accessibility in IGF2BP1, 2, and 3-KO; **B** VENN diagram for genes associated with the significant increase of chromatin accessibility in IGF2BP1, 2, and 3-KO; **C** Gene ontology analysis of biological and molecular functions for genes with significantly decreased chromatin accessibility; **D** Gene ontology analysis of biological and molecular functions for genes with significantly increased chromatin accessibility.

**Supplemental Fig. 4.**
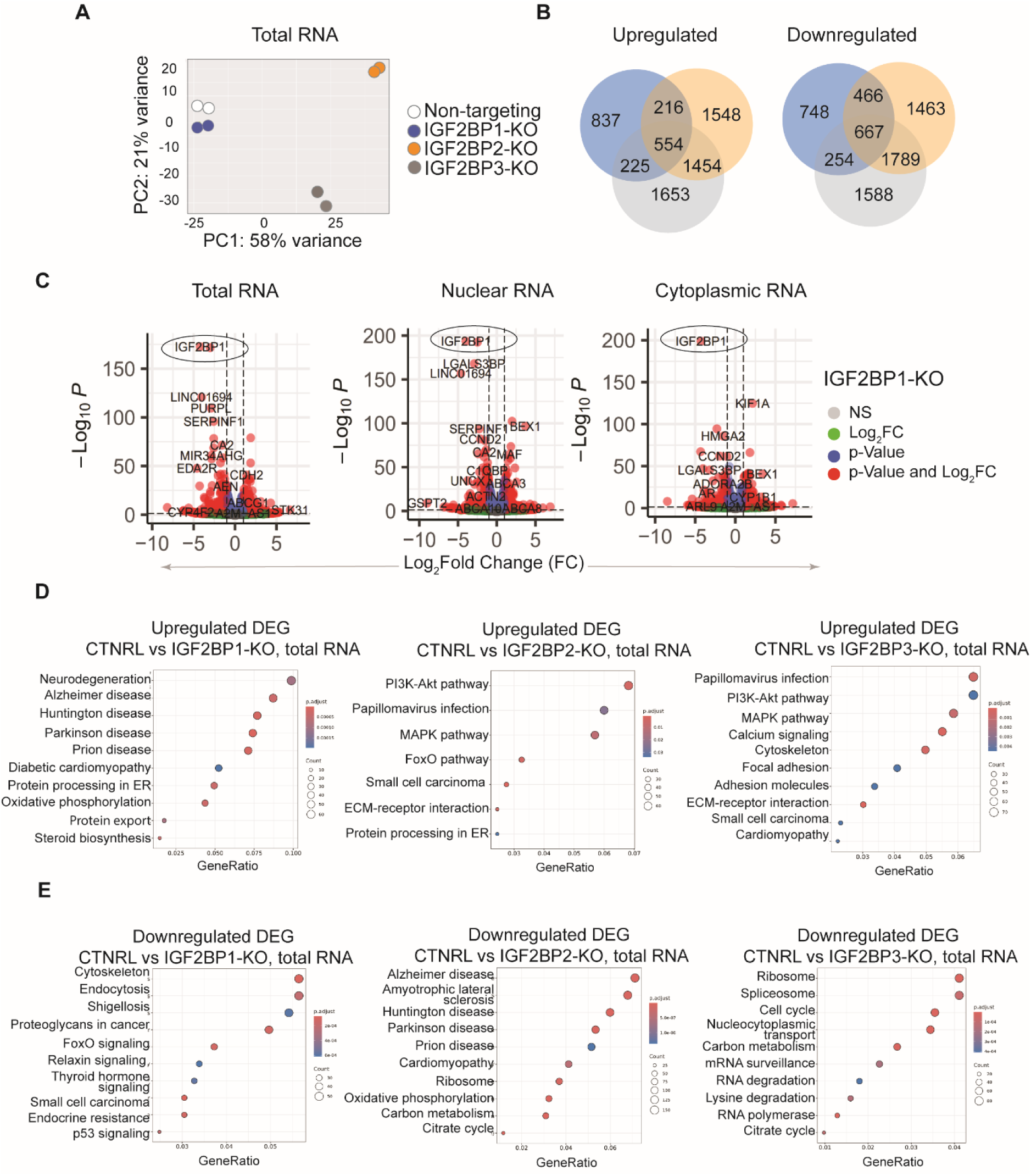
(Supplemental to main Fig.2). RNA-seq analysis of IGF2BP-KO in HEK293T cells. **A** A principal components (PC) analysis of indicated total RNA-seq samples; **B** VENN diagrams indicating commonly and uniquely upregulated (left) and downregulated (right) differentially expressed genes (DEG) between IGF2BP1, 2, and 3-KO compared to CRISPR/Cas9 control; **C** Volcano plots depicting DEG, including significantly downregulated IGF2BP1, in total, nuclear, and cytoplasmic fractions of RNA in IGF2BP1-KO cells; **D, E** KEGG enrichment for upregulated (**D**) and downregulated (**E**) differentially expressed genes (DEG) in total RNA isolated from HEK293T cells with IGF2BP-KO.

**Supplemental Fig. 5.**
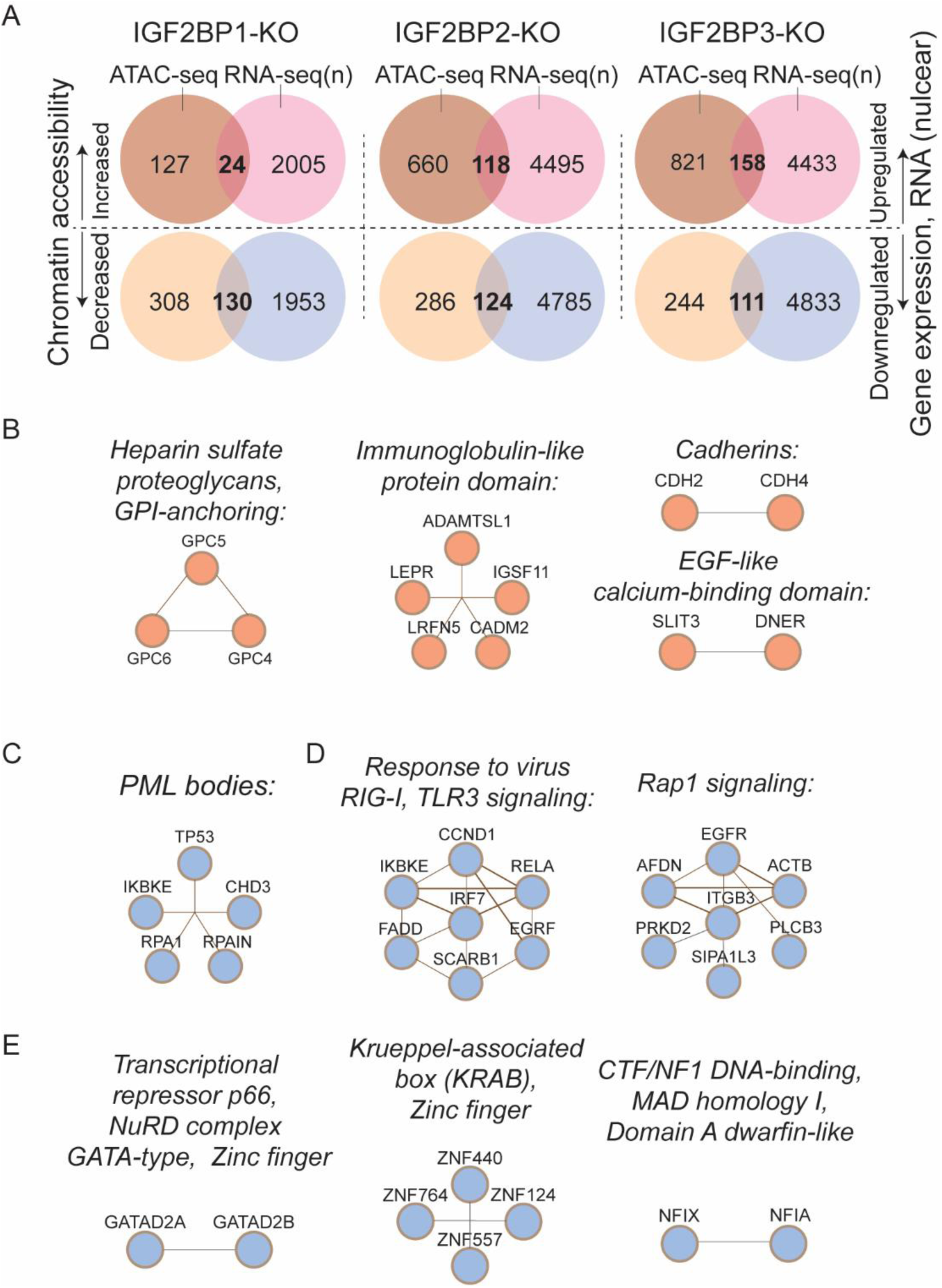
(Supplemental to main Fig.2). Integrative ATAC-seq and RNA-seq analysis for IGF2BP1-3-KO in HEK293T cells. **A** VENN diagram depicting shared increased or decreased chromatin accessibility nearby protein coding genes and corresponding upregulation or downregulation of gene’s expression; **B** Enrichments with functional protein domains in increased chromatin/upregulated mRNA fraction were common for three IGF2BP1-3-KO and included various components of plasma membrane; **C-E** Enrichments with protein domains in decreased chromatin/downregulated mRNA fractions identified functional nodes specific for IGF2BP1-KO (**C**), IGF2BP2-KO (**D**), and IGF2BP3-KO (**E**).

**Supplemental Fig. 6.**
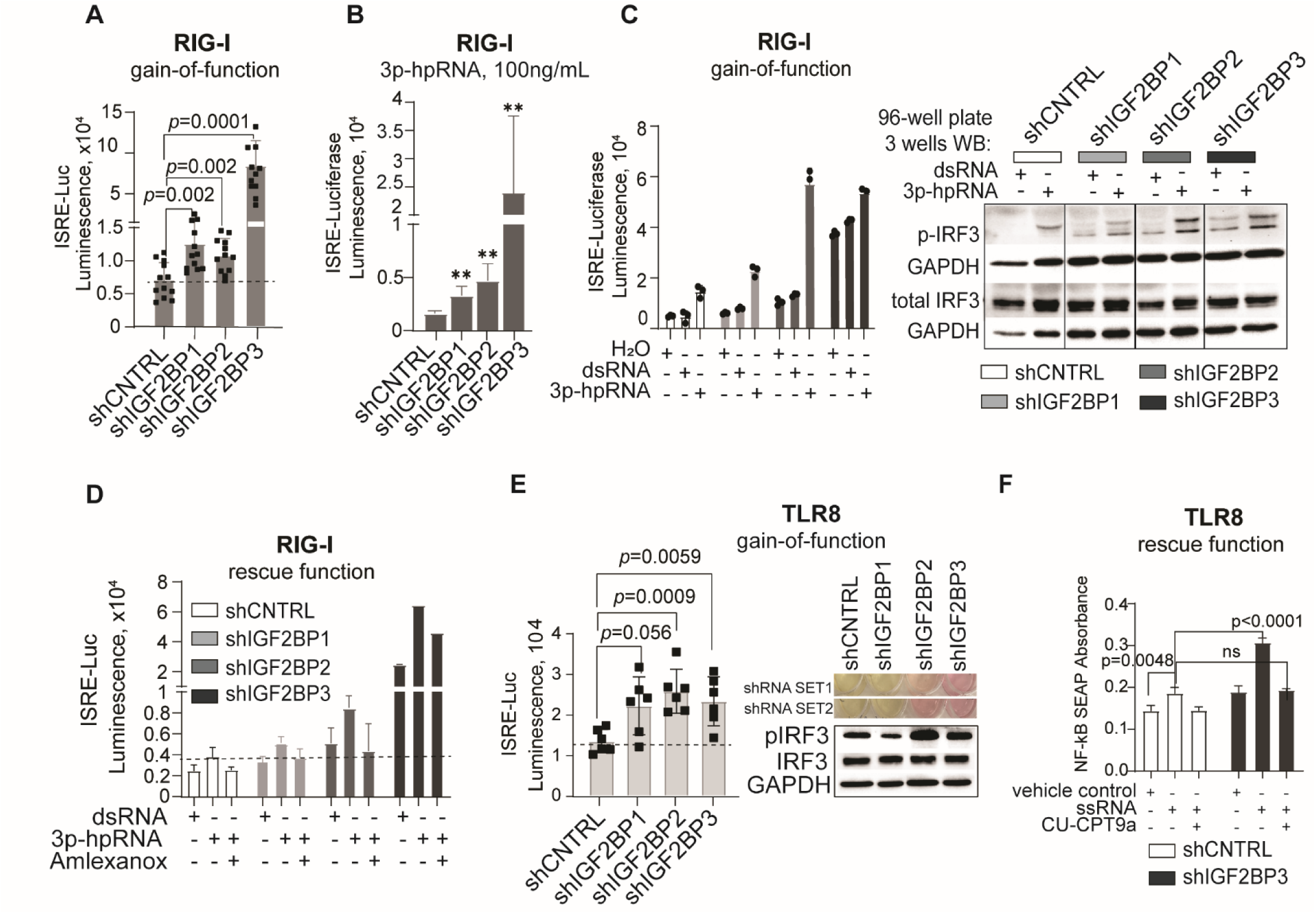
(Supplemental to main Fig.3). The effect of IGF2BPs levels on activity of RNA sensing reporter systems. **A** ISRE-driven luciferase activity in untreated HEK293T cells with RIG-I overexpression and IGF2BP1, 2, 3 genes knockdown, relative to non-targeting shRNA control. Data presented as a mean ± SD of individual values (*n*=12) of three independent biological repeats. **B** ISRE-driven luciferase activity in RIG-I and IGF2BP-KD expressing HEK293T cells treated with 100 ng/mL 3ppp-hairpin RNA (3p-hpRNA); Data are mean ± SD (*n*=4). **C** The analysis of ISRE-dependent transcription in a single experiment (*n*=3 wells of a 96-well plate): ISRE-driven luciferase in RIG-I and IGF2BP-KD expressing HEK293T cells treated with vehicle control, 100 ng/mL unphosphorylated dsRNA control or dsRNA 3p-hpRNA, n=3 wells of 96-well plate (left); western blot analysis of phosphorylated and total IRF3 proteins collected from the corresponding wells of 96-well plate (right). **D** ISRE-driven luciferase expression in RIG-I overexpressing HEK293T cells with IGF2BP-KD treated with RIG-I agonists alone (3p-hpRNA) or in combination with corresponding antagonists of TBK1/IKKε (Amlexanox). Error bars represent mean ± SD (*n*=3). **E** ISRE-driven luciferase activity in HEK293T cells overexpressing TLR8 (gain-of-function) and IGF2BPs loss-of-function (KD) without treatment and corresponding western blot of phosphorylated IRF3 and images of cultures; Data are mean ± SD of individual values (*n*=6) of three independent biological repeats. **F** NF-κB-driven SEAP levels in TLR8-OE HEK293T cells with IGF2BP3-KD treated with TLR8 agonists/ligand (ssRNA) alone or in combination with corresponding antagonists/inhibitor of TLR8 receptor (CU-CPT9a). Data is presented as mean ± SD (*n*=3).

**Supplemental Fig. 7.**
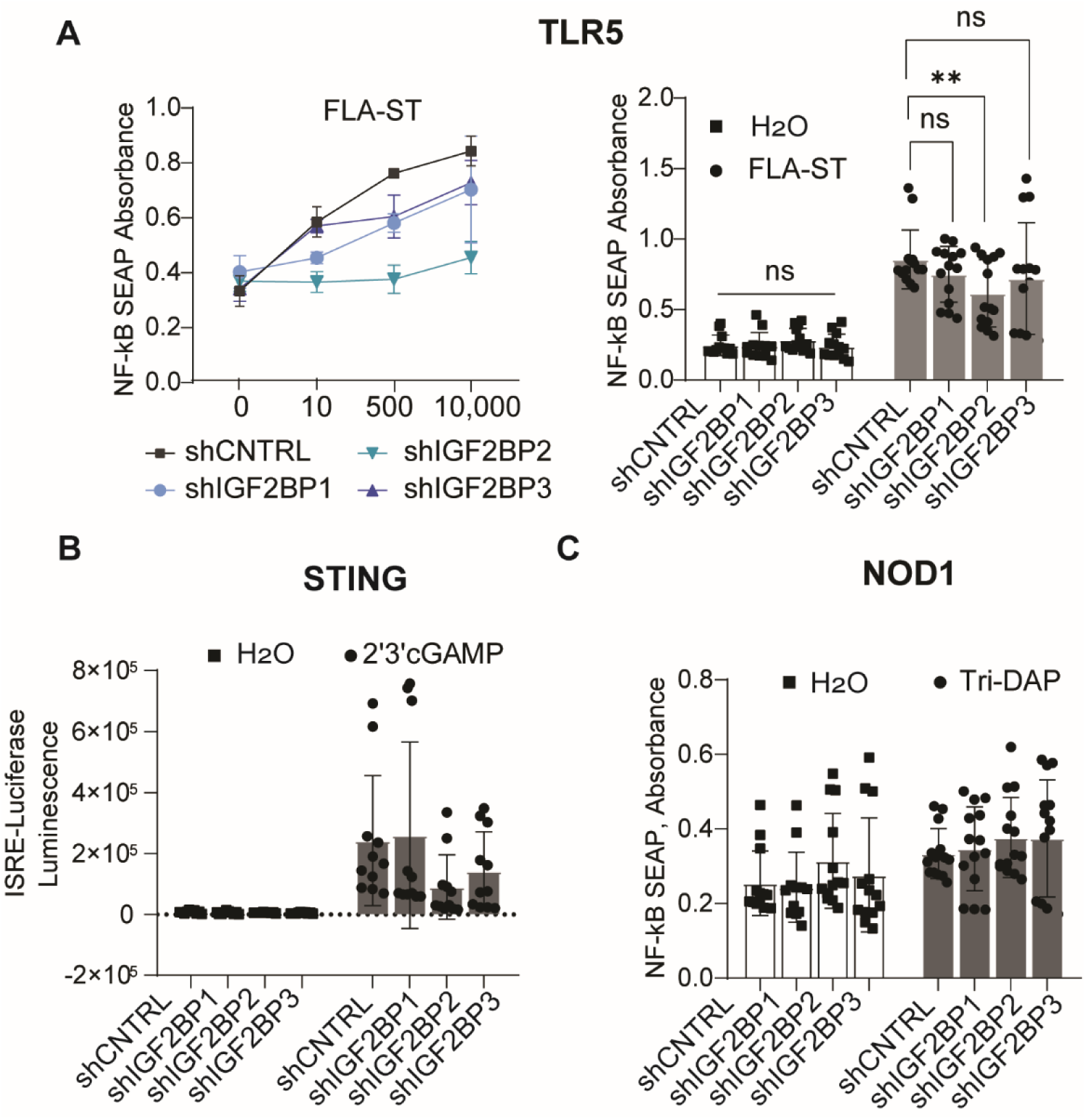
(Supplemental to main Fig.3). The effect of IGF2BPs levels on activity of non-RNA sensing reporter systems. **A** NF-kB-driven SEAP levels in HEK293T cells with IGF2BP-KD and endogenous levels of TLR5, treated with TLR5 agonist FLA-ST at 10ng/mL, 500ng/mL, 10,000 ng/mL (left); the results of three independent biological replicates of FLA-ST treatment at 10mg/mL (right). Data presented as a mean ± SD of individual values (*n*=14) from three independent repeats, unpaired two-tailed *t*-test \*\**p* < 0.01, *p*>0.05 non-significant (ns); **B** ISRE-driven secreted luciferase activity in HEK293T cells with IGF2BP-KD and endogenous levels of STING, treated with 100mg/mL 2’3’cGAMP. Data are presented as mean ± SD of individual values (*n*=11) from three independent repeats; **C** NF-kB-driven SEAP levels in HEK293T cells with IGF2BP-KD and endogenous levels of NOD1, treated with 10mg/mL of Tri-DAP. Data of three independent biological replicates is presented as mean ± SD of individual values (*n*=14).

**Supplemental Fig. 8.**
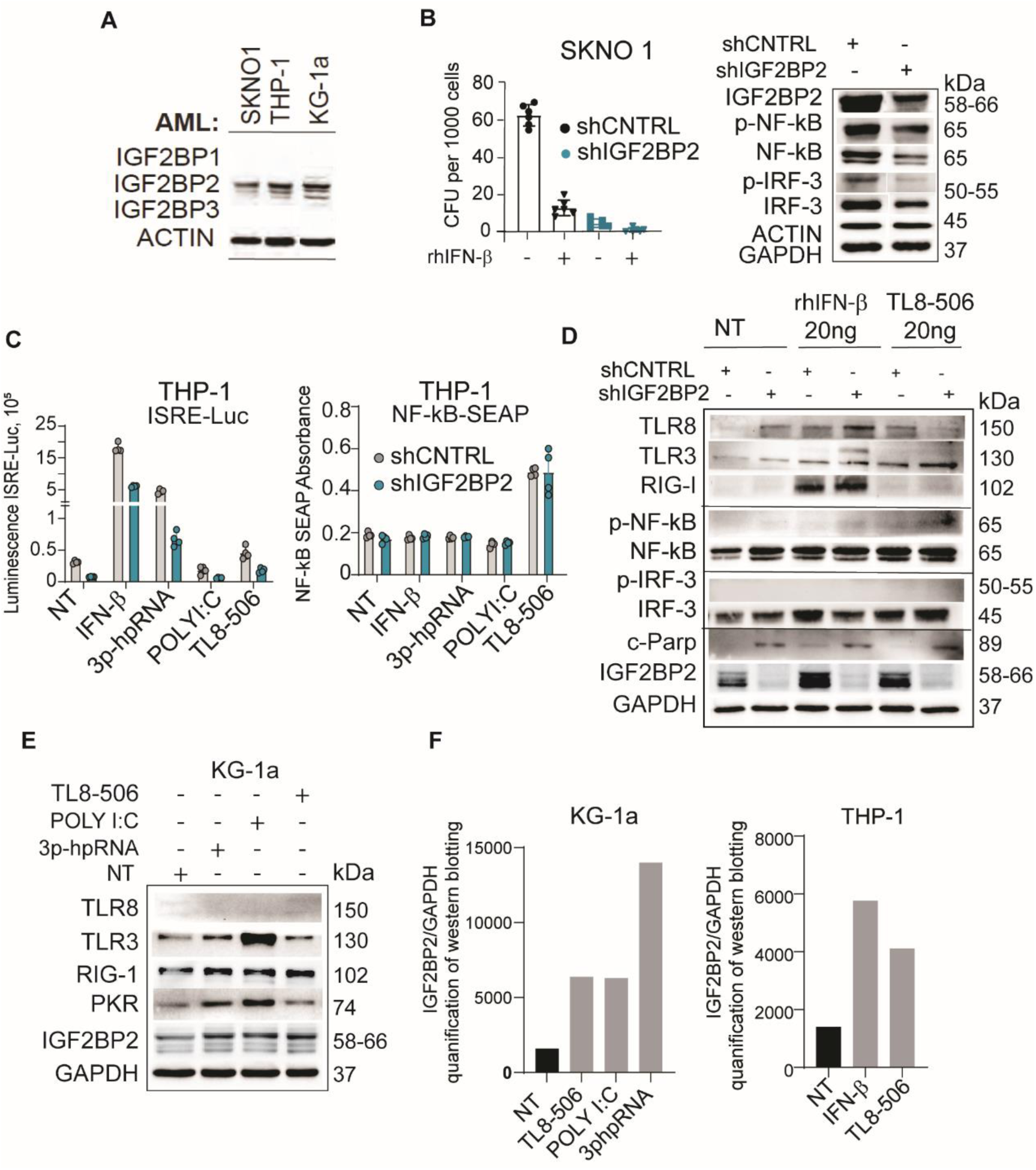
, (Supplemental to main Fig. 4). IGF2BPs expression in myeloid leukemia cells and their role in innate immune signaling. **A** IGF2BP1-3 levels in three AML cell lines assessed by western blotting. **B** Colony forming units’ assay for SKNO1 AML with IGF2BP2-KD treated as indicated, day 21-post-transduction (left); corresponding western blot analysis (right panel); Data are mean ± SD (*n*=5). **C** ISRE-driven luciferase and NF-κB-driven SEAP in THP-1 cells with IGF2BP2-KD treated with IFN-β (20 ng/mL), RIG-I ligand 3p-hpRNA (100 ng/mL), TLR3 ligand poly(I:C) (10 μg/mL), TLR8 ligand TL8-506 (20 ng/mL). Data are mean ± SD (*n*=3). **D** Western blot analysis of THP-1 treated with IFN-β (20 ng/mL) or TLR8 ligand TL8-506 (20 ng/mL) for 12 hours, compared to non-treatment control (NT). **E** Western blot analysis of KG-1a AML treated with RIG-I ligand 3p-hpRNA (100 ng/mL), TLR3 ligand poly(I:C) (10 μg/mL), TLR8 ligand TL8-506 (20 ng/mL). **F** Quantification of IGF2BP2 protein levels (IGF2BP2 relative to GAPDH expression) using ImageJ software and western blot analysis in Suppl.8E (KG-1a) and Suppl.Fig.8D. (THP-1), treated as indicated for 12 hrs (treatment conditions see Supplemental Methods, Table 6).

**Supplemental Fig. 9.**
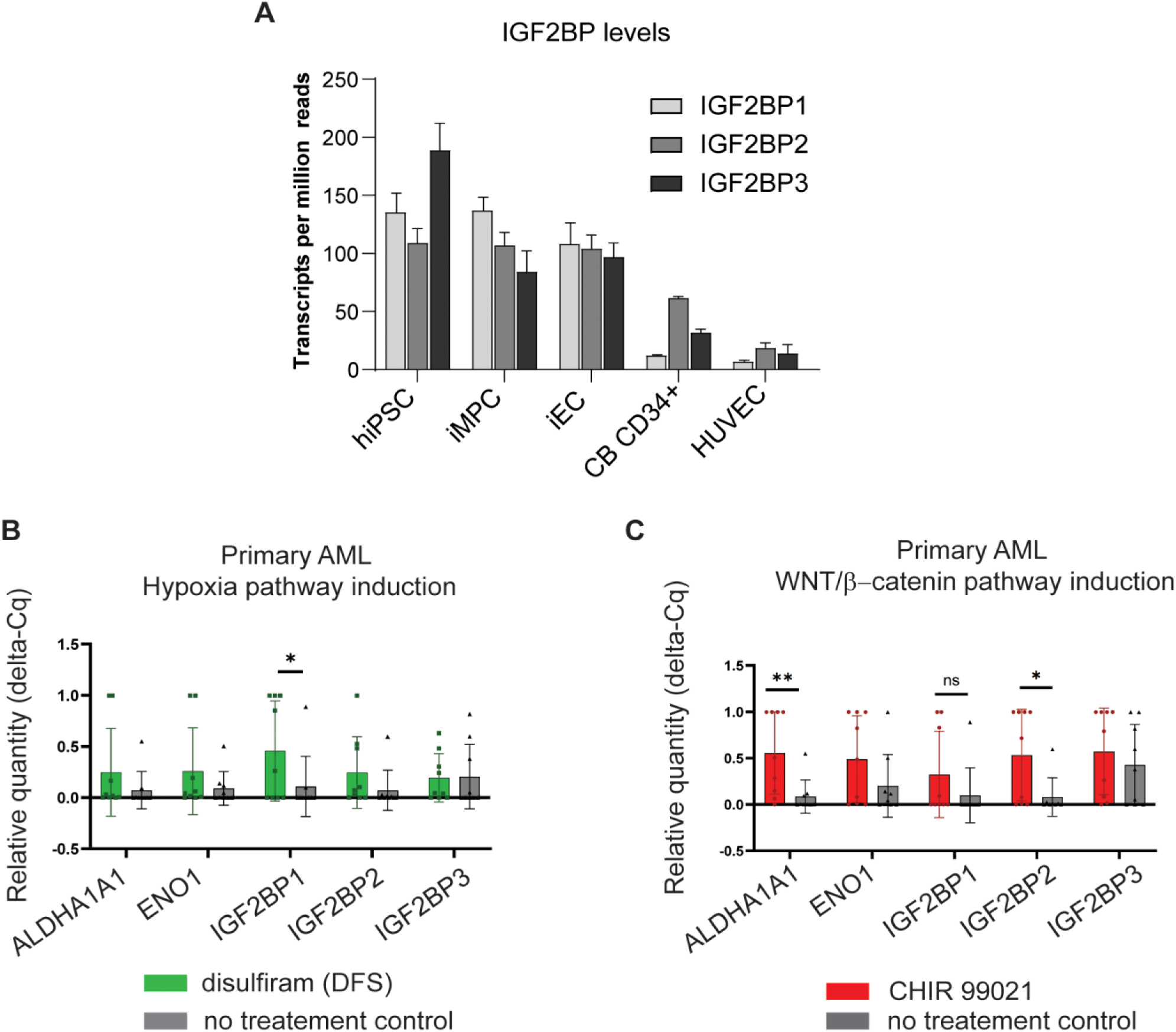
(Supplemental to main Fig.5). IGF2BP expression patterns in AML and their role in innate immune signaling. **A** IGF2BP1-3 mRNA expression levels in hiPSCs, hiPSCs-derived myeloid progenitor cells (iMPC), endothelial cells (iEC), primary human cord blood (CB) CD34+ cells and human umbilical cord vein endothelial cells (HUVEC) assessed by RNA-seq, transcripts per million reads (tpm), results are presented as a mean ±SEM (hiPSC (*n*=6), iCMP (*n*=8), iEC (*n*=5), CB CD34^+^(*n*=2), HUVEC (*n*=2)). **B, C** Cumulative qPCR analysis of ALDHA, ENO1, and IGF2BPs expression in three primary AML (#1256, 1341, 1346) with and without disulfiram (DSF) treatment (1mM for 24hrs) (**B**), or CHIR99021 treatment (10mM for 48hrs) (**C**). Data is presented as mean ±SD of individual values (*n*=9) from three independent repeats. Indicated statistical analysis is unpaired two-tailed *t*-test, \*\**p* < 0.01; \**p* < 0.05.

**Supplemental Fig. 10.**
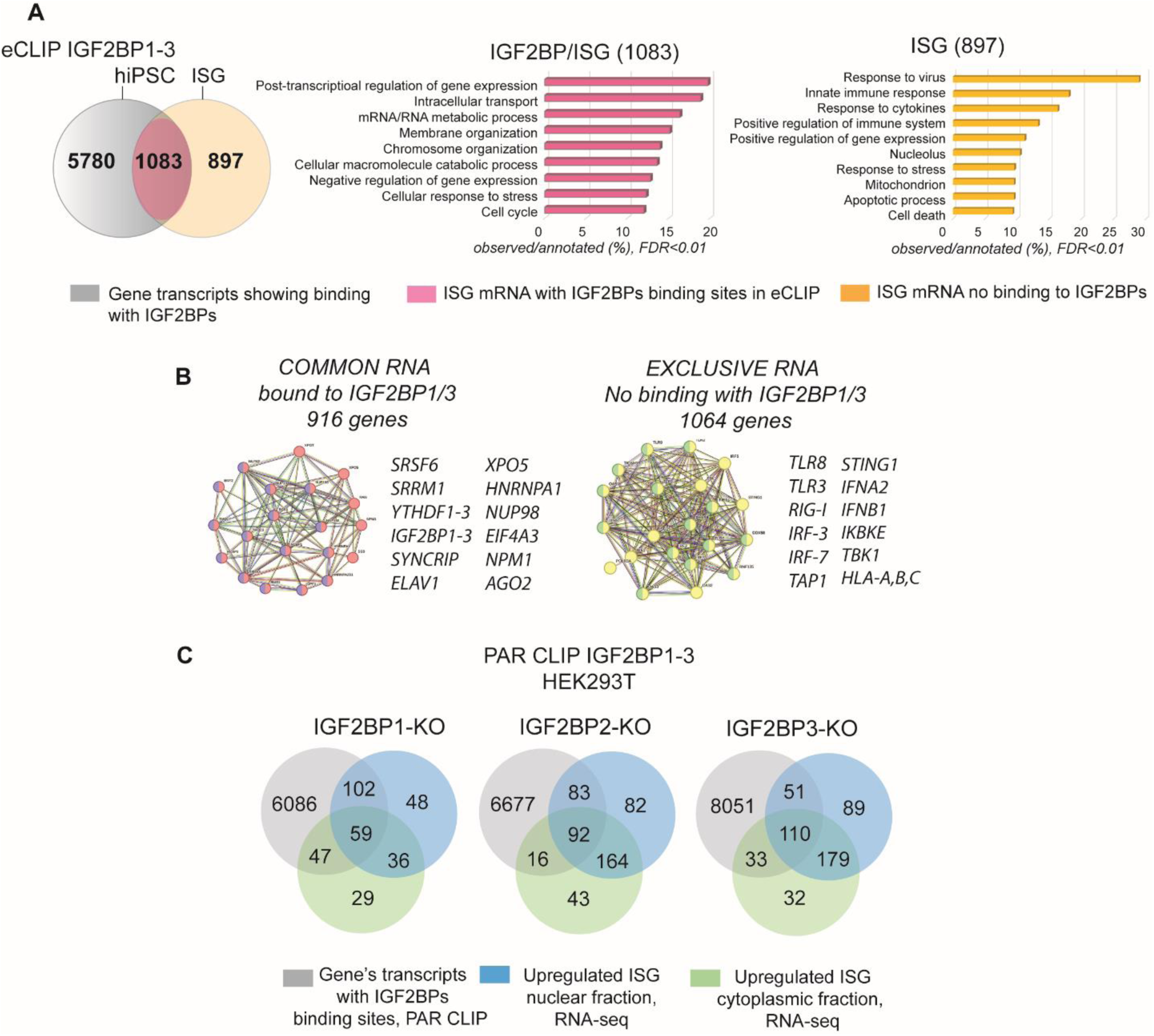
(Supplemental to main Fig.6). IGF2BPs show selective binding with interferon-stimulated gene mRNAs. **A** Visual representation of relationships (VENN diagram) between datasets generated from enhanced RNA-protein crosslinking and immunoprecipitation (eCLIP) for RNA bound to IGF2BP1, 2, 3 in hiPSC and ISGs annotated in the human gene database GeneCards; corresponding Gene Ontology (GO) pathway analysis for ISGs with (left) and without (right) IGF2BPs binding sites. **B** ISGs annotated in GeneCards database (∼2000 genes, main Fig.5A); corresponding gene cluster and pathway analysis for ISGs bound (left) and not bound to IGF2BPs (right). **C** VENN diagrams for PAR CLIP in HEK293T cells and upregulated cytoplasmic or nuclear ISGs, isolated from individual IGF2BP-KO in HEK293T cells (analyzed in this study, Fig. 2G).

